# Fiber-based Probes for Electrophysiology, Photometry, Optical and Electrical Stimulation, Drug Delivery, and Fast-Scan Cyclic Voltammetry In Vivo

**DOI:** 10.1101/2024.06.07.598004

**Authors:** Nicolette Driscoll, Marc-Joseph Antonini, Taylor M. Cannon, Pema Maretich, Greatness Olaitan, Valerie Doan Phi Van, Keisuke Nagao, Atharva Sahasrabudhe, Emmanuel Vargas, Sydney Hunt, Melissa Hummel, Sanju Mupparaju, Alan Jasanoff, Jill Venton, Polina Anikeeva

## Abstract

Recording and modulation of neuronal activity enables the study of brain function in health and disease. While translational neuroscience relies on electrical recording and modulation techniques, mechanistic studies in rodent models leverage genetic precision of optical methods, such as optogenetics and imaging of fluorescent indicators. In addition to electrical signal transduction, neurons produce and receive diverse chemical signals which motivate tools to probe and modulate neurochemistry. Although the past decade has delivered a wealth of technologies for electrophysiology, optogenetics, chemical sensing, and optical recording, combining these modalities within a single platform remains challenging. This work leverages materials selection and convergence fiber drawing to permit neural recording, electrical stimulation, optogenetics, fiber photometry, drug and gene delivery, and voltammetric recording of neurotransmitters within individual fibers. Composed of polymers and non-magnetic carbon-based conductors, these fibers are compatible with magnetic resonance imaging, enabling concurrent stimulation and whole-brain monitoring. Their utility is demonstrated in studies of the mesolimbic reward pathway by simultaneously interfacing with the ventral tegmental area and nucleus accumbens in mice and characterizing the neurophysiological effects of a stimulant drug. This study highlights the potential of these fibers to probe electrical, optical, and chemical signaling across multiple brain regions in both mechanistic and translational studies.

## 1. Introduction

Since their conception in the early 1900s, electrical recording and stimulation of neural activity have laid the foundation to study brain circuits.^[1]^ More recently, advances in genetic and optical tools have empowered neuroscientists to probe neural pathways with cell-type specificity, uncovering mechanisms underlying behavior and physiology in health and disease.^[2,3]^ Optogenetics using microbial rhodopsins permits excitation and silencing of specific neurons with millisecond precision.^[4]^ Similarly, genetically encoded calcium and neurotransmitter indicators enable imaging and photometric recording of neurochemical signaling with cell-type specificity in behaving subjects.^[5–7]^ Despite their utility in rodent models, genetic techniques remain challenging in non-rodent species and require extensive regulatory review in human patients. Complementing clinically viable electrical recording and stimulation, fast-scan cyclic voltammetry (FSCV) can measure the release of redox-active neurotransmitters, such as dopamine (DA).^[8]^

Tools capable of electrical recording, stimulation, FSCV, optogenetics, photometry, and drug and gene delivery would not only enable mechanistic studies of brain circuits in rodent models but could also empower correlative experiments between precise genetic approaches and translational alternatives. This further highlights the need to pair neural interfaces with other preclinical techniques such as whole-brain activity mapping with functional magnetic resonance imaging (fMRI).^[9,10]^ Although optical methods reliant on passive glass or polymer waveguides are compatible with MRI,^[11]^ metallic electrodes often introduce magnetic field inhomogeneities, reducing the signal-to-noise ratio (SNR) and spatial resolution during functional imaging.^[12]^

Here we sought to create MRI-compatible probes suitable for behavioral experiments that are simultaneously capable of bidirectional electrical, optical, and chemical recording and modulation of neuronal signaling via genetic and translational techniques. Integrating these functions is challenging in any materials platform, including mature wafer-bound semiconductors, and fabrication without consideration for tissue mechanics can yield devices with limited biocompatibility and long-term utility. Consequently, the development of integrated tools benefits not only from miniaturization but also the use of low-modulus materials.^[13]^

We applied thermal drawing to seamlessly integrate low magnetic susceptibility and high-conductivity carbon-based electrodes, low-loss polymer optical waveguides, and microfluidic channels into miniature and flexible fibers. Although prior work on multifunctional fiber-based probes has demonstrated integration of metal (tin, indium, tungsten, stainless steel, and copper) and carbon-composite (carbon-loaded polyethylene or polycarbonate) electrodes, these materials had either a limited charge injection capacity (CIC) or limited compatibility with magnetic resonance imaging.^[14–17]^ Similarly, prior work on multi-material fibers employed polymer waveguides which exhibited optical power losses in the range of 1.5 – 2.7 dB/cm,^[14,15]^ which were acceptable for optogenetic neuromodulation. Photometric recording, however, requires low-loss optical transmission and is typically performed with commercial silica fibers, which suffer from poor mechanical compatibility with neural tissue.^[15,18–22]^

Here, we employ 20 µm carbon nanotube (CNT) yarn electrodes that not only demonstrate exceptional CIC and low impedance for electrical stimulation and electrophysiological recording, but also have magnetic susceptibility (χ_m_ = –10 ppm) suitable for MRI.^[23]^ Furthermore, the high surface area of CNT electrodes facilitates superior adhesion of DA as compared to FSCV field-standard carbon fiber microelectrodes, thereby increasing the sensitivity of FSCV measurements.^[24]^ Further, we integrate waveguides with a PMMA (poly(methyl methacrylate), refractive index n_PMMA_ = 1.50) core,^[25]^ an industry standard for low-loss transmission, and a low-index THVP cladding (terpolymer of tetrafluoroethylene, hexafluoropropylene, and vinylidene fluoride, n_THVP_ = 1.35).^[26]^ These innovations in electrode and waveguide materials choices are coupled with engineering solutions to enable facile back-end interfaces with multifunctional fibers which are produced in a single convergence-drawing step from macroscale models (preforms).

The resulting MRI-compatible Polymer-based Optical-electrical-chemical neuroLogical Interface (POLI) fibers integrate electrophysiology, chemical sensing, optogenetics, photometry, and fluid delivery capabilities within footprints smaller than those of silica fibers used for photometry and optogenetics. We demonstrate that the POLI fibers can be applied to simultaneous interrogation of multiple signaling modalities (neural activity, calcium signaling, and DA release) across two structures of the mesolimbic pathway, a circuit critical for reward and motivation processing.^[27]^

## 2. Results

### 2.1. POLI Fiber Design and Fabrication

The POLI fiber probe was designed with six 20 µm-diameter CNT yarn electrodes and a 200 µm-diameter PMMA/THVP optical waveguide embedded within an insulating polycarbonate (PC) cladding. A microfluidic channel (40×100 µm^2^) composed of a PC wall was separated from the cladding via a thin layer of styrene-ethylene-butylene-styrene (SEBS) elastomer (**Figure 1a**). The constituent polymers were selected for the relative similarity of their glass transition temperatures (80-185 °C, **Supporting Table S1**) to enable thermal co-drawing.^[28–30]^ Additionally, PMMA and THVP were selected for their low absorption and high refractive index contrast in the visible range. The SEBS layer surrounding the microfluidic channel was introduced to facilitate connections of each fiber element to its respective back-end interface. Due to its relatively weak adhesion to PC, SEBS enables peeling of the microfluidic channel from the side of the fiber for in-line fluidic connection to external tubing, and isolating the optical and electrical components for connectorization (**Supporting Figure S1**).^[31,32]^

**Figure 1.**
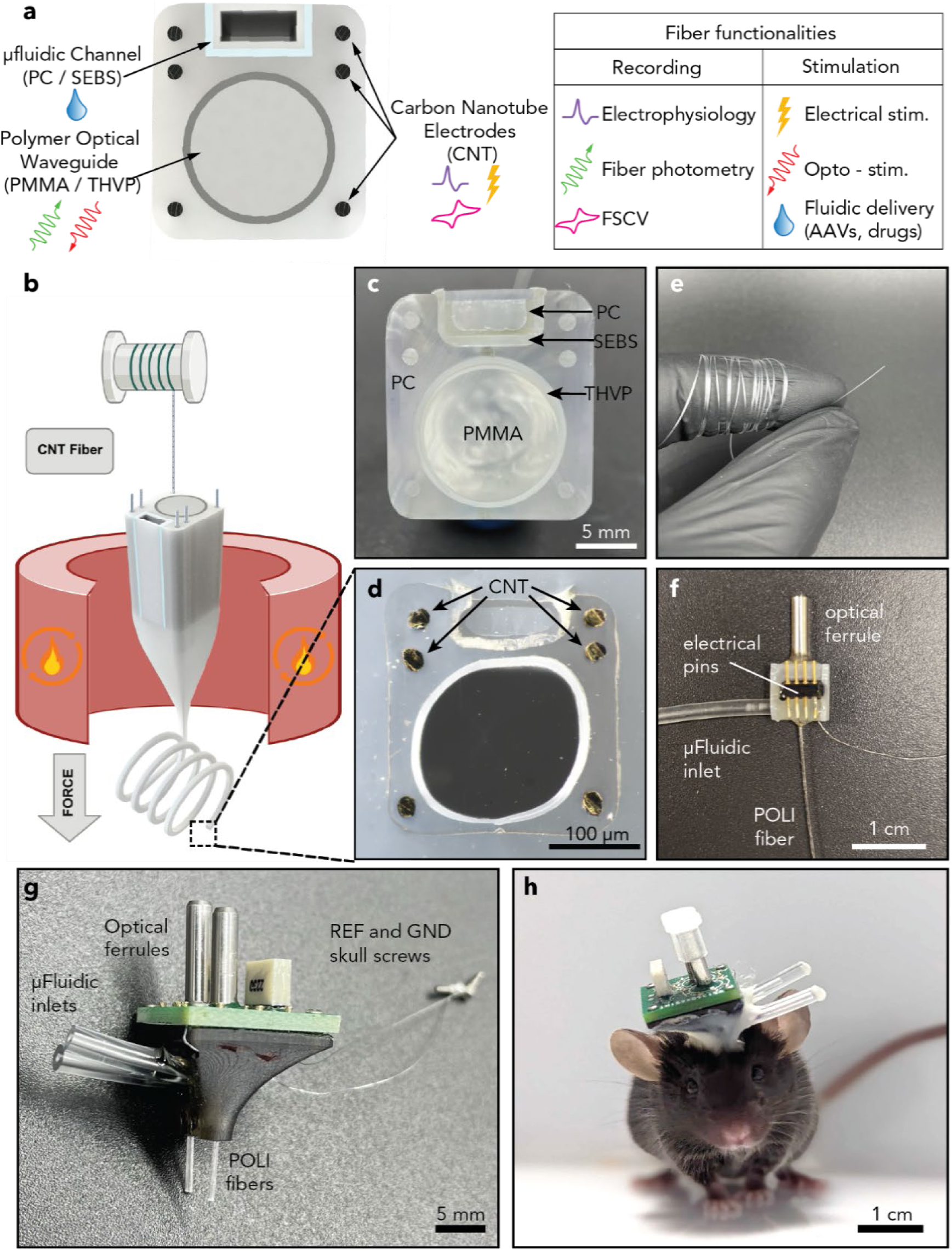
Fabrication of the POLI fiber. (a) schematic of the cross-section of the POLI fiber, highlighting its recording and stimulating capabilities. (b) Diagram of the convergence drawing process used to fabricate the POLI fiber. (c) Photograph of the macroscale preform highlighting the polymer combination choice and the channels used for electrodes, and microfluidics (scale bar = 5 mm). (d) Optical micrograph of the fiber cross-section highlighting the cross-sectional geometry and embedded CNT microwires (scale bar = 100 μm). (e) Photograph highlighting the flexibility and microscopic size of the POLI fiber microfluidics. (f) A fully assembled POLI fiber with electrical pins, optical ferrule and microfluidic inlet (scale bar = 1 cm). (g) Photograph of a dual-implant assembly that leverages PCB and Omnetics connector to facilitate connectorization. This implant simplifies the implantation process of fiber devices targeting the VTA and NAc (scale bar = 5 mm). (h) Photograph of a freely moving mouse with the dual-fiber implant (scale bar = 1 cm).

To fabricate the fiber, a macroscale preform (1.99×2.20 cm^2^ cross-section, 22.9 cm long) containing PC, PMMA, THVP, and SEBS components was produced through machining and consolidation techniques, and channels were drilled into the PC body to accommodate CNT yarn. The preform was then thermally drawn at a temperature of 260-280 °C, and the 20 µm diameter CNT yarn was converged from spools into the open channels of the preform to form the electrodes (**Figure 1b, c**). The drawing procedure delivered flexible fiber with cross-sectional dimensions of 306 ± 17 µm by 342 ± 8 µm (**Figure 1d, e**). Minor distortion of the PMMA/THVP optical waveguide was observed but did not significantly impact transmission losses of the PMMA core as discussed below. To facilitate interfacing with electrical recording and stimulation equipment, fiber-coupled light sources, and tubing leading to micropumps, sections of fiber were outfitted with backend interfaces comprising a printed circuit board (PCB), fluidic tubing, and optical ferrules in a 3D-printed housing to permit attachment of either one (**Figure 1e**) or two (**Figure 1f**) POLI fibers. The PCBs bore a 16-channel Omnetics® electrical connector, and openings allowed for 2 optical ferrules to be threaded through the PCB. In each device realization, each individual fiber remained capable of performing all six functions. The complete assemblies weighed 2.292 g and were compatible with stereotaxic brain surgery in mice (**Figure 1h**).

### 2.2. POLI Fiber Characterization

The functional performance of each POLI fiber element was first assessed independently ex vivo. To quantify the ability of the waveguides with PMMA (n_PMMA_ = 1.4956, λ = 488 nm) core and THVP (n_THVP_ = 1.35, λ = 589 nm) cladding to collect and transmit optical signals (**Figure 2a**), we measured their loss coefficients (α, dB/cm, **Figure 2b**) and numerical apertures (NA, **Figure 2c.i-d.iii**). Previously reported multifunctional fiber-based probes relied on waveguides with PC core (n_PC_ = 1.60, T_g_ =130 - 170 °C) and cyclic olefin copolymer (COC, n_COC_ = 1.55, T_g_ = 158°C) cladding.^[33]^ While suitable for optogenetics, these waveguides were too lossy to enable photometric recordings. Thus, we evaluated the loss coefficients and NA of polymer waveguides composed of several combinations of core and cladding: PMMA/THVP, PC/PMMA, and COC/PMMA with similar core-cladding diameters (400 µm / 420 µm). Consistent with the high refractive index contrast between PMMA and THVP and lower absorption of PMMA as compared to PC in the visible range, we found the largest NA = 0.53±0.04 and lowest loss coefficient α = 0.55±0.14 dB/cm (n = 3 samples) for PMMA/THVP fibers as compared to COC/PMMA (NA = 0.28±0.05, α = 1.5±0.39 dB/cm) and PC/PMMA (NA = 0.44±0.03, α = 1.10±0.25 dB/cm) waveguides, respectively (**Figure 2b, c**). Note that PMMA/THVP fibers exhibit higher NA than commercial silica fibers with a similar core diameter (400 µm, manufacturer NA = 0.5, measured NA = 0.50±0.022), which yields improved light collection for fiber photometry. Due to minor deviations from the cylindrical shape in custom-drawn fibers, experimentally measured NA values were slightly lower (17-27%, **Supporting Figure S2**) than those calculated from the reported refractive indices, but followed anticipated trends. Although polymer waveguides have greater loss coefficients than silica fibers, the resulting attenuation is minimal at rodent-brain scales (6.1% for 5 mm) and is likely acceptable for studies in larger organisms such as non-human primates (58.8% for 7 cm, **Figure 2a**).^[34]^ Additionally, the substantially lower stiffness of polymer-based fibers compared to silica fibers is anticipated to improve mechanical compatibility with brain tissue.

**Figure 2.**
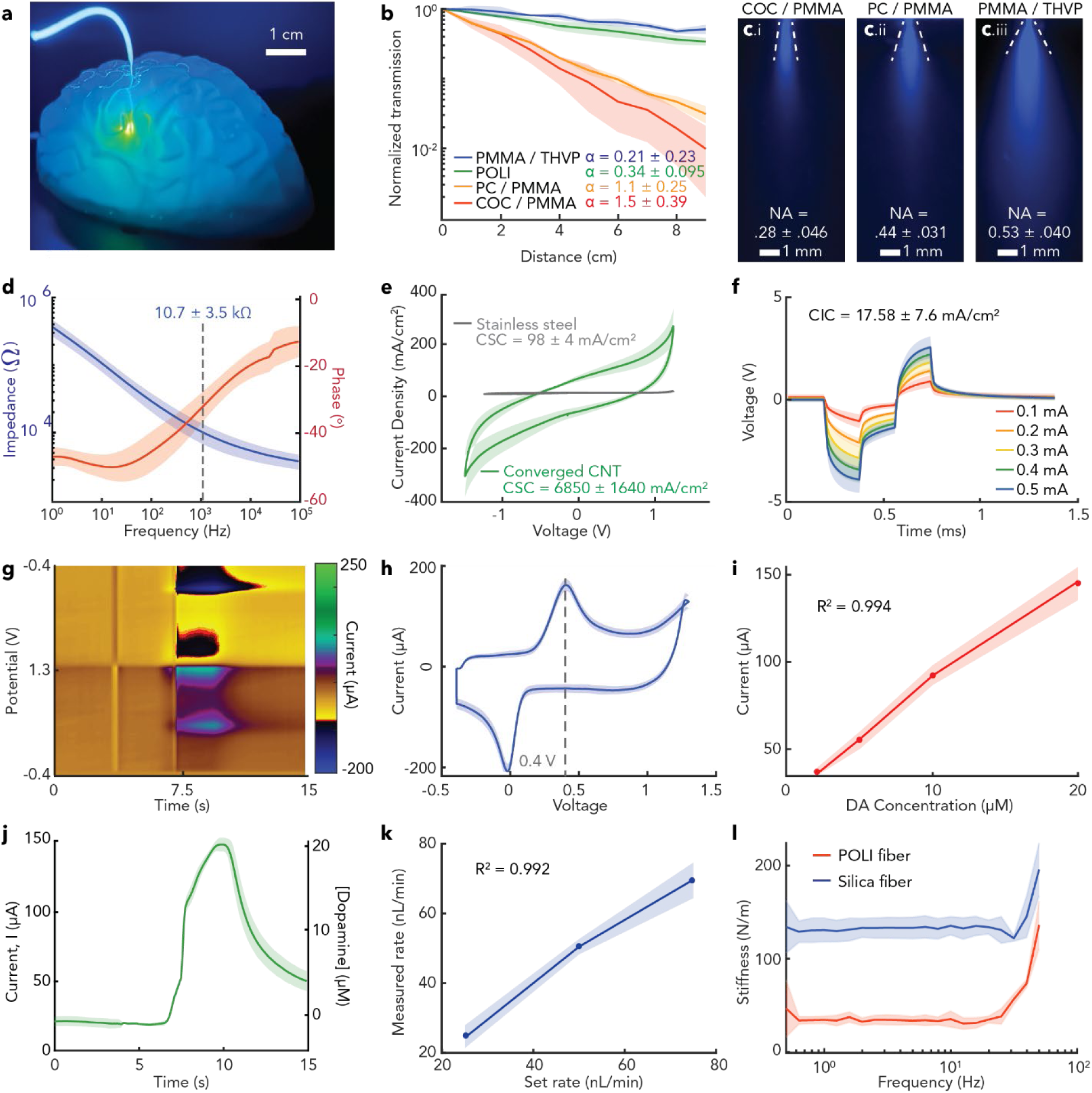
Benchtop characterization of the POLI fiber. (a) Picture of a PMMA/THVP waveguide implanted in a fluorescein-doped agarose brain transmitting blue (470 nm) light to elicit a fluorescent response (∼520 nm). Blue light is visible along the patch cord due to high input optical power, which permitted visual observation of the fluorescent response; the power of optical leakage along the fiber due to impurities or microscale defects is orders of magnitude below the power of light transmitted to the fiber tip, and as observed, did not elicit a fluorescent response along the length of the fiber (scale bar = 1 cm). (b) Evaluation of optical loss for 400 µm PC/PMMA, COC/PMMA, and PMMA/THVP polymer fibers, as well as the 200 µm PMMA/THVP optical waveguide of the POLI Fiber. Optical loss described as mean decibel loss ± s.e.m. (n = 3 samples). (c) Numerical aperture measurement of 400 µm PC/PMMA, COC/PMMA, and PMMA/THVP polymer fibers, expressed as mean ± s.e.m. (n = 3 samples, scale bar = 1 mm). (d) Bode plot of impedance magnitude and phase of the POLI fiber-embedded 20 µm-diameter CNT microelectrodes (n = 3 fibers, 18 electrodes total). (e) Cyclic voltammograms of the CNT electrodes and same-size comparison stainless-steel electrodes in phosphate-buffered saline (PBS) and their respective cathodic charge storage capacity values. (f) Voltage transient response of CNT electrodes to symmetric, biphasic, charge-balanced, current pulses of 250 µsec half phase and a 250 µsec interphase delay, and corresponding cathodic charge injection capacity. (g) Fast scan cyclic voltammetry response of a CNT electrode to a bolus of 20 µM DA solution delivered at t = 7 s recorded in a flow cell. (h) A representative voltammogram of CNT electrode response to 20 µM DA solution, showing DA oxidation peak at +0.4 V and reduction peak at 0 V. (i) DA calibration curve demonstrating a linear relationship between detected current and dopamine concentration in solution (R^2 = 0.994, n = 3). (j) Current measured at +0.4 V vs. time for n = 5 trials of 20 µM DA solution bolus delivered in the flow cell, converted to DA concentration using the calibration curve shown in (i). (k) Measured rate of fluid injection via POLI fiber microfluidic channel vs. rate set on the infusion pump. Infusions were delivered at set rates between 25, 50, and 75 nl/min and are represented as mean ± s.e.m. (n = 3). (l) Stiffness measured via dynamic mechanical analysis of POLI fibers (n = 3) compared to a similarly sized silica fiber across the frequency ranges of locomotion, respiration, and heart rate.

We next evaluated the electrochemical properties of the 20 µm-diameter CNT electrodes embedded within the POLI fiber. Electrochemical impedance spectroscopy (EIS) revealed characteristic capacitive behavior (**Figure 2d**), with a mean |Z| value of 10.7±3.5 kΩ at 1 kHz (n = 3 fibers, 18 electrodes) which was only slightly affected by repeated stimulation pulsing (n = 5, 100,000 pulses) with a slight decrease of impedance observed after the first 2000 cycles due to electrode conditioning, beyond which the impedance remained stable. (**Supporting Figure S3**)

CNT electrodes possess a wide electrochemical water window of –1.5 to +1.2 V vs. Ag/AgCl; within this window, electrical stimulation pulses can be delivered without inducing hydrolysis. We measured the cathodic charge storage capacity (CSCc) of the CNT electrodes in this window to be 6850±1636 mC/cm^2^, which is significantly greater than same-size stainless steel electrodes characterized over the same voltage range (98.0±4.1 mC/cm^2^) (**Figure 2e**). The CSC of CNT electrodes is significantly larger than that of standard clinical stimulation electrodes (platinum, CSC = 0.55 mC/cm^2^; iridium oxide, CSC = 4 mC/cm^2^)^[35,36]^. Note that the CSC value measured here for the CNT electrodes exceeds that of previous reports of CNT-based electrodes.^[37]^ We hypothesize that this measured CSC can be attributed to the large electrochemically active surface area of the rough CNT yarn electrodes, which the geometric surface area used in our CSC calculations did not account for. The charge injection capacity (CIC) of the CNT electrodes was measured via voltage transient testing with biphasic current pulses, increasing in amplitude until the maximum cathodic excursion potential (E_mc_) crossed the water window limit of –1.5 V. The CNT electrodes in the POLI fiber exhibit a CIC of 17.6±7.6 mC/cm^2^, which is substantially higher than that of stainless steel (CIC = 1.5±0.5 mC/cm^2^) as well as clinical alternatives (platinum, CIC = 0.35 mC/cm^2^; iridium oxide, CIC = 0.87 mC/cm^2^)^[36,38]^, indicating a greater amount of charge that can be safely injected during a stimulation pulse (**Figure 2f**). Notably, platinum or platinum-iridium electrodes could be integrated within thermally drawn polymer fibers via the same convergence process used here, however CNTs were chosen for the POLI fiber due to their charge injection characteristics and MRI compatibility.

The low impedance and wide electrochemical stability window (–1.5 - 1.2 V) of CNT electrodes also make them suitable for FCSV.^[8,39–41]^ FSCV is commonly performed using brittle carbon fiber electrodes, whereas CNT yarn offers flexibility, robustness, and greater sensitivity while retaining electrochemical stability.^[42,43]^ The FSCV capabilities of CNT electrodes within POLI fibers were first assessed in vitro in a flow cell with solutions of DA in HClO_4_ diluted to concentrations of 2 μM - 20 μM in PBS using a standard DA waveform scanning between –0.4 -1.3V at a rate of 100 V/s and sampling frequency of 10 Hz against Ag/AgCl reference electrodes (**Figure 2g,j,i-m**). A color plot illustrates the current response of the CNT electrodes to 10 µM DA in solution (**Figure 2g**), and a corresponding cyclic voltammogram reveals known DA redox peaks (**Figure 2h**). The peak current at the oxidation voltage (0.4 V) varies linearly over a range of physiological DA concentrations (**Figure 2i,j**, R^2^ = 0.994, n = 3 fibers), with a limit of detection (LOD) of 0.1916 µM and a limit of quantification (LOQ) of 0.581 µM, suggesting the utility of CNT electrodes for DA concentration measurements in vivo.^[44]^

The microfluidic channels within POLI fibers (40 × 100 µm^2^) were assessed by recording the rate of fluid passage driven by a syringe pump. Consistency (1:1) between the pump and the fiber flow rates was found across the microfluidic channels of multiple POLI fibers (**Figure 2k**, R^2^ = 0.992, n = 3 devices).

The stiffness of POLI fibers (360-380 x 400-420 µm^2^, n = 3 devices) was assessed via dynamic mechanical analysis (DMA) over a range of frequencies corresponding to heartbeat, respiration, and locomotion (**Figure 2l**). Compared to a silica waveguide (diameter = 400µm, n = 3 fibers), the POLI fiber exhibited a significantly lower bending stiffness in single cantilever mode, which was comparable to previously reported polymer-based multifunctional fibers with long-term tissue stability.

### 2.3. POLI Fiber Functional Characterization In Vivo

We first evaluated the abilities of POLI fibers to perform photometric recordings of activity indicators, electrical stimulation (including during MRI), and FSCV measurements of DA. Photometric recordings with PMMA/THVP fiber platform were first compared to those performed with commercial 400 µm diameter silica multimode fibers, the most common photometry platform. PMMA/THVP fibers thermally drawn to 200 µm or 400 µm diameter (core/cladding ratio=95%) were implanted into the contralateral whisker sensory cortices (S1BF) of Thy1-GCaMP6s mice broadly expressing a fluorescent calcium indicator GCaMP6s in excitatory neurons (**Supporting Figure S4a**).^[45]^ In these mice, silica fibers were implanted into the same brain region in the opposite hemisphere (**Supporting Figure S4b**). Fluorescence increases in response to whisker flicks were recorded by both 200 µm (n = 8 trials, 1 female mouse) and 400 µm (n = 15 trials, 3 mice - 2 male and 1 female **Supporting Figure S7. Immunohistochemical evaluation of chronically bilaterally implanted silica and POLI fibers.** Immunofluorescent quantification of astrocytic (GFAP) and microglial (Iba1, CD68) markers surrounding a 300-400µm section of POLI-fiber and a 400 μm silica fiber implanted into the NAc of male DAT^IRES*cre*^ mice (N=4) (HC PLAPO CS2 10x/0.40 Dry objective, WLL: 85% Power, Speed: 400Hz, 405: Intensity (3.80), Gain (33.76); 499: Intensity (0.5), Gain (27.62); 554: Intensity (3.66), Gain (25.49), 653: Intensity (2), Gain (33.76), scale bar = 100 µm). Average fluorescent area and average fluorescence intensity of Iba1 (a-d), GFAP (e-h), and CD68 (i-l) as well as representative confocal images at the implant tips of POLI fiber (-1.25ML, +1.2AP, -4.3DV) and silica fiber (+1.25ML, +1.2AP, -4.3DV) 1 month post implantations. GFAP average mean intensity; p = 0.0421, two-sample t-test. Data are presented as mean values +/- s.d.) PMMA/THVP fibers with an SNR comparable to that of the 400 µm silica fiber (n = 15 trials, 6 mice - 3 females and 3 males) (**Supporting Figure S4c-h**).

Given that the diameters of PMMA/THVP waveguides within POLI fibers are 200 µm, suitable for photometric recordings, we then applied these devices to test the ability of the CNT electrodes to drive neural activity and evoke calcium influxes. POLI fibers were implanted in the S1BF of Thy1-GCaMP6s mice, and the embedded CNT electrodes were used to deliver bipolar electrical stimulation at frequencies of 10 Hz (**Figure 3a-c**) and 130 Hz (**Figure 3d-e**). These stimulation frequencies and currents varying between 35-215 µA are typical for therapeutic neuromodulation.^[46]^ Robust increases in normalized GCaMP6s fluorescence (ΔF/F_0_) were recorded in response to 10 Hz stimulation above 88 µA, and the threshold was lower for 130 Hz stimulation indicating a dose-dependent response. These observations corroborate the ability of POLI fibers to simultaneously deliver electrical stimulation and photometrically record evoked neuronal activity.

**Figure 3.**
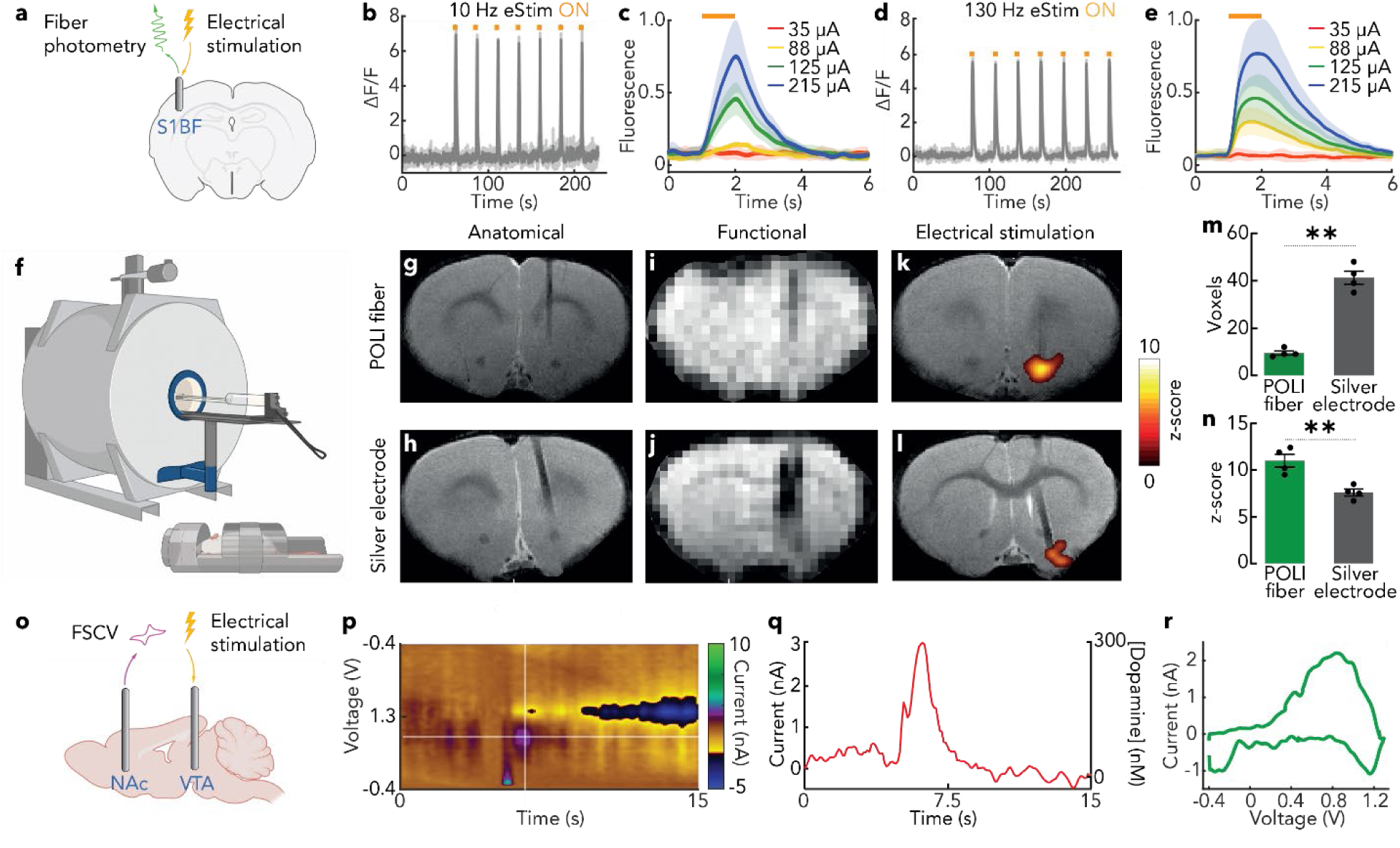
*In vivo* validation of POLI fiber functions. (a) Schematic of the experiment. The POLI fiber was implanted in the whisker sensory cortices (S1BF) of Thy1-GCaMP6s mice, and used to concurrently stimulate electrically through the embedded CNT electrodes and record stimulation evoked calcium influxes using fiber photometry. (b-e) Calcium transient was measured following an electrical stimulation train of 10 Hz (b,c) and 130 Hz (d,e), with current varying between 35-215 µA. Fluorescence represented as a mean fluorescence (ΔF/F_0_) ± s.e.m, with (c) and (e) fluorescence normalized to the max fluorescence value. (f) High field MRI acquisition was performed with a 20 cm-bore 9.4 T Bruker small animal scanner with a custom-made 30 mm single surface coil used as a transceiver. (g-l) Anatomical scans were acquired using a T2-weighted rapid acquisition with refocused echoes (RARE) pulse sequence. Functional scans were performed using T2 *-weighted EPI sequence for detection of stimulus-induced BOLD contrast. Deep brain stimulation of 0.1mA for 2s at a frequency of 60Hz was applied preceded by 10s baseline scan and followed by 48s recovery scan, repeated over 30 cycles. (n = 8 female Sprague-Dawley rats) (m,n) Preprocessing of the functional scans was performed and scans were aligned to high-resolution anatomical images; maximum signal amplitude during an interval of 6s after stimulation onset was measured and compared to the average preceding baseline interval via Student’s t-test to evaluate statistical significance of z-scores. To quantify voxel count, a threshold of SNR< 5 was applied on slices in which the implant was observed. ** corresponds to P<0.01. (o) FSCV performed through POLI fiber was used to detect DA release in NAc in response to electrical stimulation in VTA supplied via an electrode, with a representative color plot (n = 1 Sprague-Dawley rat) (p) and voltage transient (q) shown. DA concentration was calculated from maximum at its oxidation peak in a representative voltammogram (n = 6 stimulations) (r).

We then applied POLI fibers to deliver electrical stimulation in a preclinical context where changes in blood-oxygen-level-dependent (BOLD) signal recorded during MRI are used to assess the effects of deep brain stimulation (DBS). POLI fibers were implanted in a nucleus accumbens (NAc), a subcortical structure of the basal ganglia implicated in reward processing, in Sprague Dawley rats (n = 8). These experiments were performed in parallel in rats implanted with a pair of silver electrodes (356 µm in diameter) traditionally used for DBS during MRI in rodents (**Figure 3f**). The 6-electrode POLI fibers reduced the tissue damage associated with the insertion of two separate electrodes (**Figure 3g-l**). Anatomical scans were acquired using a T2-weighted rapid acquisition with refocused echoes (RARE) pulse sequence with 18 1-mm thick slices with the field of view (FOV) of 20 mm × 20 mm, the echo time (TE) of 34.7 ms,^[47]^ and the repetition time (TR) of 2 s. Functional scans were performed using a T2*-weighted echo-planar imaging (EPI) sequence for detection of stimulus-induced BOLD contrast, with TE of 16 ms, TR of 2 s, FOV of 20 mm × 20 mm, and image size of 40 mm × 40 mm.^[48]^ The DBS (60 Hz, 100 µA, 2 s ON, 48 s OFF, repeated for 30 cycles) applied via the CNT electrodes within POLI fibers resulted in greater increase in BOLD signal than identical stimulation protocol delivered via silver electrodes under continuous EPI scans (POLI fiber 11.0 ± 1.4, silver electrodes: 7.6 ± 0.7, n = 4 rats/group, **Figure 3m,n**). This is likely due to the masking of voxels with high BOLD signals adjacent to the silver electrodes, with the POLI fiber mitigating susceptibility-induced artifacts.^[49]^

To test the utility of POLI fibers for FSCV recordings of DA, these probes and the two 230 µm bipolar stainless-steel electrodes (MS303, Plastics One) were implanted into the NAc core and the ventral tegmental area (VTA) of a Sprague Dawley rat (**Figure 3o**). The VTA is comprised of ∼ 56% DA neurons in rats, and the release of DA from NAc-projecting VTA DA neurons is the hallmark of reward.^[50,51]^ Following 24 bi-phasic electrical pulses (2 ms, 300 μA) delivered in the VTA, we observed robust phasic DA concentration release in the NAc core (**Figure 3p-r**). The DA concentration was extrapolated via a post-experiment electrode calibration procedure (Experimental Section). Evoked DA levels were consistent over 6 stimulation epochs separated by 5-min intervals with an average current of 3.07 ± 0.32 nA, corresponding to a concentration of 199 ± 20.55 nM, which is comparable to prior reports. ^[52,53]^

### 2.4. POLI Fiber as a Tool to Assess Drug-Induced Perturbations in the Reward Circuit

Maladaptive changes at key nodes of mesolimbic reward circuit, the VTA and NAc, accompany chronic drug and alcohol use.^[54–56]^ Developing tools to reveal changes in the VTA and NAc signaling in response to drugs of abuse holds the potential to advance the study and treatment of substance use disorders. Here, we evaluated the utility of POLI fibers to interrogate the DA projection circuit between the VTA and the NAc at baseline conditions as well as in the presence of cocaine, a well-characterized modulator of DA signaling in the brain.^[57]^ Adult transgenic DAT::Cre mice were implanted with POLI fibers in the VTA and the NAc simultaneously (n = 10, 5 male, 5 female). A fluorescent dopamine indicator dLight1.1 under pan-neuronal promoter human synapsin (hSyn) and packaged into an adeno-associated virus (AAV9) was delivered into the NAc via the microfluidic channel of the POLI fiber during implantation. The excitatory opsin ChrimsonR fused to a fluorescent protein mScarlet under CamkIIa promoter and packaged into an AAV9 vector was delivered into the VTA (**Figure 4a**).^[7,56]^ Following 14 days of incubation, experiments consisting of electrophysiological recordings in the NAc and VTA, photometric recording of dLight1.1 fluorescence in the NAc, optogenetic stimulation of DA neurons in the VTA, and electrical stimulation in the VTA or NAc were performed.

**Figure 4.**
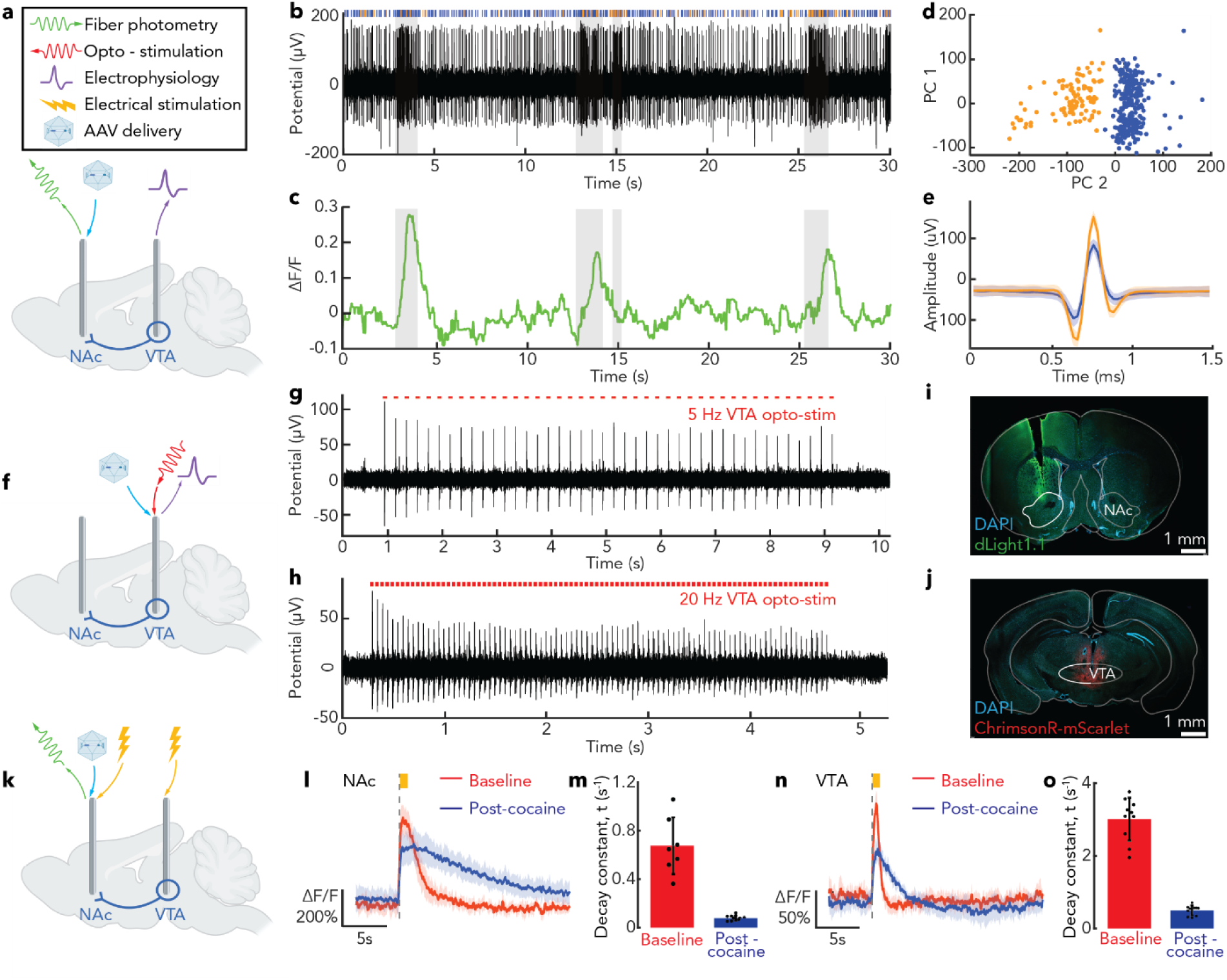
Probing mesolimbic DA dynamics and drug-induced perturbations with POLI fiber. (a) Mice were implanted with the dual implant device, comprised of two POLI fibers targeting the NAc and the VTA, and transfected with DA fluorescent indicator dLight1.1 into the NAc via AAV delivery through the POLI fiber fluidic channel. (b) Endogenous electrophysiology activity recorded in VTA, showing multiunit neural activity with phasic and tonic firing. Blue dots indicate spikes detected for sorting. (c) Fiber photometry recording of endogenous DA dynamics in NAc concomitant with recordings in (b). Shaded areas in (b) and (c) indicate phasic firing interspersed with tonic firing in unshaded areas (see **Supporting Figure S5**). Large DA transients observed in NAc photometry recordings are time-locked to bursts of tonic firing of the neurons in VTA. (d) Principal components analysis and clustering of neural spike waveforms recorded in VTA, and (e) mean spike waveforms for the two neuronal units. (f) Mice were implanted with the same dual implant POLI fiber device and transfected with a red excitatory opsin ChrimsonR in the VTA via AAV9 delivery through the POLI fiber channel to enable simultaneous optogenetic stimulation and electrophysiology recording. (g,h) Optogenetic stimulation-evoked activity of VTA neurons with (g) 5 Hz stimulation and (h) 20 Hz stimulation. (i,j) Fluorescent images show expression of (i) dLight1.1 in the NAc and (j) ChrimsonR in the VTA (HC PLAPO CS2 10x/0.40 Dry objective, WLL: 85% Power, Speed: 400Hz, 405: Intensity (2.00), Gain (12.54); 499: Intensity (2.00), Gain (22.30); 554: Intensity (2.00), Gain (10.86), scale bar = 1mm). In the same cohort of animals used in (a-e), electrical stimulation was applied either to the VTA or to the NAc while recording DA dynamics in NAc via fiber photometry. (l) Electrical stimulation of DA axon terminals in the NAc produced robust DA transients recorded via photometry. Stimulation epochs were recorded before and after intraperitoneal (IP) administration of cocaine (20 mg/kg, n = 5 epochs for each animal). Data are presented as mean ± s.e.m. (m) Electrical stimulation of cell bodies in the VTA produced robust DA transients in the NAc, though lower in amplitude compared to NAc stimulation. Stimulation epochs were recorded before and after cocaine administration (20 mg/kg, IP, n = 5 epochs for each animal). Data are presented as mean ± s.e.m. (n,o) Following administration of cocaine, stimulation-evoked dLight1.1 transients exhibited a significantly slower decay compared to the pre-cocaine transients due to inhibition of DA reuptake.

First, we recorded spontaneous activity, including electrophysiology in the VTA and NAc with simultaneous photometry recording of DA dynamics via dLight1.1 photometry in the NAc (n = 10 mice, 5 male, 5 female). **Figures 4b** and **4c** show example simultaneous electrophysiological recordings in the VTA and dLight1.1 photometry readings in the NAc. Principal components analysis and k-means clustering of the data in **Figure 4b** revealed the spike waveforms of two putative neuronal units (**Figure 4d,e**). These VTA units exhibited tonic firing with an average rate of 7.46 Hz, interspersed with bouts of high-frequency (25.2 Hz) phasic activity (**Figure 4d, Supporting Figure S5**) which is characteristic of VTA DA neurons. Epochs of phasic firing in these VTA neurons preceded dLight1.1 fluorescence transients recorded in the NAc (**Figure 4b-c**), further suggesting their DA-producing identity. Examples of additional recordings are shown in Supporting Figure S6.

Next, we recorded electrophysiological signals in the VTA while delivering 635 nm laser pulses (5Hz and 20Hz frequency; 10 ms pulse width) through the waveguide in the same POLI fiber to drive DA neurons expressing ChrimsonR (**Figure 4f**). Both 5 Hz and 20 Hz optogenetic stimulation drove activity of the VTA DA neurons (**Figure 4g,h**). Expression of ChrimsonR-mScarlet in the VTA and dLight1.1 in the NAc were corroborated with confocal microscopy in postmortem brain samples (**Figure 4i,j**).

Further, we assessed the biocompatibility of POLI fibers against that of comparably-sized silica fibers (Supporting Figure S7) at a time point of one-month post-implantation in the NAc of DAT::Cre mice using a standard immunohistochemical panel to identify glial scarring (glial fibrillary acidic protein, GFAP), macrophage activation (ionized calcium-binding adapter molecule 1, Iba1) and microglia activation (cluster differentiation 68, CD68). The POLI fiber immune response was found to be lower or comparable to that of the silica fibers across all inflammatory markers, a consistent finding with their experimentally measured lower bending stiffness (Fig. 2l), supporting their ability to perform several diverse modalities of stimulation and recording without introducing additional tissue damage.

Cocaine is known to inhibit DA reuptake within the mesolimbic pathway.^[53,58,59]^ To determine whether cocaine-induced changes in DA dynamics could be resolved using recording and stimulation capabilities of the POLI fibers, these probes or Notch fibers – functionally equivalent prototypes of POLI fibers (**Supplemental Note 1**) were implanted into the NAc and VTA of C57BL/6J wild-type mice (n = 5, male), with dLight1.1 delivered to the NAc as described above (**Figure 4k**). Using the CNT electrodes in POLI fibers, neurons in the VTA were electrically stimulated (60 Hz; 2 ms biphasic pulses; 200 µA; 0.5 s ON, 30 s OFF) and dLight1.1 signals corresponding to stimulation-evoked DA were measured in the NAc, where DA neurons in the VTA send their projections, before and after cocaine administration (20 mg/kg, intraperitoneal) (**Figure 4k**). We observed a slower decay of stimulation-evoked dLight1.1 transients following cocaine injection, indicating inhibited DA reuptake in the NAc (**Figure 4m**). This experiment was repeated using Notch fibers, with electrical stimulation this time applied directly to the axon terminals in NAc rather than to the cell bodies in the VTA. The stimulation-evoked DA transients recorded in the NAc were larger for direct NAc stimulation as compared to VTA stimulation, and cocaine administration caused a similar slowing of DA decay by inhibiting DA reuptake (**Figure 4l**). In these experiments, we showed that fiber-integrated CNT electrodes could be used for electrical stimulation to evoke DA release in the NAc, either by stimulating the cell bodies in the VTA, or by directly stimulating the axon terminals in the NAc. Furthermore, integrated optical waveguide could concomitantly record the DA dynamics via dLight1.1 photometry. Consistent with prior work, following administration of cocaine, we observed a decrease in a rate of decay of stimulation-evoked dLight1.1 fluorescence stemming from the blocking of the dopamine transporter (DAT) (**Figure 4n,o**).^[60–63]^

## 3. Conclusion

Through materials optimization and convergence thermal drawing, we have engineered MRI-compatible POLI fibers capable of bidirectional optical, electrical, and chemical interrogation of neuronal brain circuits in vivo. The lower losses and higher NA of PMMA/THVP waveguides within POLI fibers, with respect to previously reported polymer waveguides, permitted photometric recording of calcium and dopamine indicator fluorescence with SNR comparable to commercial silica fibers. Low impedance, high CIC, and high CSC of the CNT yarn electrodes permitted their application to electrophysiology, electrical stimulation, and FSCV-detection of DA in vivo. Furthermore, POLI fibers enabled low-artifact functional MRI recordings allowing for monitoring of whole-brain effects of DBS. Owing to their flexibility and small footprint, POLI fibers could be safely implanted into two brain regions, which allowed interrogation of VTA-NAc DA circuits via electrophysiological and photometric approaches. Our recordings revealed changes in DA release and reuptake dynamics in the presence of cocaine, a known DA reuptake inhibitor, illustrating the potential of POLI fibers as a versatile platform for multi-site interrogations of neural circuits. By combining electrophysiological, DBS, optogenetics, fluid delivery, photometry, and FSCV capabilities in a miniature flexible fiber device, the POLI fiber is poised to advance both fundamental and preclinical neuroscience studies.

## 4 Experimental Section

### POLI Fiber Fabrication

Multifunctional fibers with six carbon nanotube microelectrodes (20 µm diameter), one 200 µm diameter PMMA/THVP polymer optical waveguide, and one rectangular microfluidic channel (100 µm × 40 µm lumen) (Fig. 1d) were produced via a thermal fiber drawing process (Fig. 1a). Thermal drawing begins with fabricating a macroscale fiber preform. In this case, we milled polycarbonate (PC) bars (McMaster-Carr, Impact Resistant Polycarbonate) with a computerized CNC Mill to form the bulk structure of the preform with six 1.59 mm electrode channels, a 14.29 mm opening for the waveguide, and a 10 mm × 5 mm rectangular channel for the microfluidic channel. Separate PC parts were consolidated in a hot press with PTFE spacers used to maintain the openings for the electrode channels, optical waveguide, and fluidic channel (185°C, 5 psi, 60 min). The optical waveguide preform was fabricated by hot-pressing THVP pellets (3M Dyneon, THVP 2030GZ) into a 200 µm-thick film (160°C, 5 psi pressure, 60 min). This THVP film was then rolled tightly around a 12.7 mm diameter PMMA rod (US Plastic Corp., Clear Extruded Acrylic Rod) to a final diameter of 14.3 mm. The waveguide preform was inserted into the central opening in the main PC preform, and they were consolidated in a vacuum oven at 160°C for 40 min. The microfluidic channel preform was produced by milling polycarbonate sheets and consolidating them into a 3.55 × 8.52 mm rectangle in the hot press using an aluminum mold and Teflon spacer placed in the lumen (185°C, 5 psi, 30 min). This PC rectangle was then hot-pressed into a layer of Styrene-Ethylene-Butylene-Styrene (SEBS, Kraton G1657) in an aluminum mold to outer dimensions of 4.97 × 9.94 mm (130°C, 5 psi, 10 min). The PC/SEBS microfluidic preform was then placed into the rectangular channel on the side of the main PC preform and consolidated in the hot press (130°C, 5 psi, 10 min). As the final step to complete the multifunctional fiber preform, the corners of the preform were rounded with a 3.55 mm rounded corner end mill.

During the fiber drawing process, the preform was placed into a vertical cylindrical oven heated to 280 °C. The carbon nanotube electrodes were incorporated into the fiber using convergence, as described in prior work^[15,64]^. Six spools of 20 µm diameter single-filament CNT fiber (Dexmat, Galvorn Fiber) were oriented above the thermal drawing oven and the ends of the CNT fibers were fed into the electrical channels of the preform. As the preform is heated above the glass transition temperature, T_g_, of the constituent polymers, it begins to flow downward and a neck forms, reducing the diameter of the preform. When the lower end of the preform reaches a capstan situated below the oven, the lower end of the preform is cut off and the necking part of the fiber is fed into the capstan. The capstan speed, *v*_*capstan*_, is slowly increased as the preform is fed into the oven at *v*_*feed*_. The ratio between *v*_*capstan*_ and *v*_*feed*_ determines the reduction factor, or the ratio between the cross-sectional area of the preform and the fiber. Here, we used *v*_*ffeed*_ = 0.25 mm/min and *v*_*capstan*_ = 1.26 m/min for a reduction factor of 1:71 between the fiber and preform dimensions. As the fiber diameter decreased to the final target size of 280 × 310 µm, the electrode channels in the preform decreased to the size of the CNT wire until they tightly constricted around the wires and pulled them into the fiber at the same rate that the fiber was drawn at.

### Polymer optical fibers fabrication

Three polymer optical fibers (PC/PMMA, COC/PMMA, PMMA/THVP) were fabricated using the thermal drawing process, using a process similar as the one outlined above. Briefly, the PC/PMMA optical waveguide was fabricated by rolling 0.05 mm PMMA films (GoodFellow, #ME301050) around a 12.7 mm PC rod (McMaster-Carr, #8571K14) until the assembly reached a diameter of 13.3 mm, which was then consolidated under vacuum at a temperature of 170°C for 40 min. The COC/PMMA optical waveguide was fabricated by first molding COC 6013 pellets (Ajedium) in a vacuum oven at 280°C for 12 hours into a 25.4 mm square bar, which was then lathed to form a 16.4 mm rod. Following, 0.05 mm PMMA films (GoodFellow) was rolled until the assembly reached a diameter of 17.1 mm, before getting consolidated in a vacuum oven at a temperature of 170°C for 40 min. Finally, the PMMA/THVP preform was fabricate by rolling 200 µm-thick THVP film (3M Dyneon THVP 2030GZ, processed as described above), around a 19.05 mm PMMA rod (US Plastic Corp. Clear Extruded Acrylic Rod) until it reached 19.8 mm in diameter, before being consolidated under vacuum at a temperature of 160°C for 40 min. Each preform was transformed into a fiber using the thermal drawing process with a draw temperature of 260°C (PC/PMMA), 215 °C (COC/PMMA), 180 °C (PMMA/THVP) until each fiber diameter reached 400 µm.

### Dual Fiber Device Assembly

Sections of the multifunctional fiber were assembled into a custom dual-fiber probe device, designed with precise spacing between the probes to enable simultaneous implantation into the mouse nucleus accumbens (NAc) and ventral tegmental area (VTA). First, the fiber was cut into 4 cm sections. The microfluidic channel was then connected to poly(vinyl chloride) (PVC) tubing (Tygon PVC, McMaster-Carr #8349T11) by peeling the first ∼20 mm of the microfluidic from the side of the fiber, trimming it to ∼3 mm in length, inserting this into the PVC tubing, and sealing the junction with UV-curable epoxy (Norland Optical Adhesive 68). Next, the PC cladding surrounding the optical waveguide and microelectrode wires was chemically etched from the upper portion of the fiber, where the microfluidic channel had already been separated, by submerging into dichloromethane (DCM, Sigma Aldrich L090000) for 2 min. The etch-exposed CNT microwires were separated from the exposed portion of the polymer optical waveguide, and the waveguide was coated with 5-minute epoxy (Devcon 14240), inserted into a 2.5 mm diameter stainless steel optical ferrule (Thorlabs #SF270-10), and allowed to cure. Excess length of the polymer optical waveguide protruding out from the top of the ferrule was trimmed, then the ferrule was polished using AlOx lapping films (Thorlabs LF5P, LF3P, LF1P, LF03P). Next, two devices were inserted into a custom-designed printed circuit board (PCB, PCBWay) by inserting the optical ferrules through precisely spaced VIAs in the PCB. The ferrules were secured in place in the PCB with UV-curable epoxy. Next, each CNT microwire from the two fiber probes was fed through a plated through-hole VIA in the board and connected with an electrode interface board (EIB) gold connector pin (Open Ephys OEPS-7010). Each of the plated through-holes was routed to a surface-mounted Omnetics electrical connector (Omnetics A79042). Finally, a custom-designed 3D-printed casing was attached to the bottom of the PCB to enclose all connections, with the two POLI fibers protruding from the bottom. The two POLI fibers were trimmed to the appropriate lengths with the help of a 3D-printed cutting mold to reach the target brain regions.

### Optical Waveguide Characterization

The optical loss coefficient of the PMMA/THVP polymer optical waveguide in the POLI fibers was measured using the cut-back method. Briefly, 12 cm sections of each fiber waveguide were cut and connected to stainless steel optical ferrules as described above. After polishing the ferrule termination, the fiber was connected via optical patch cord to a 470 nm fiber-coupled LED light source (Thorlabs M470F4) driven by an LED driver (Thorlabs LEDD1B) at a constant drive current. The light power emitted from the tip of the fiber was measured using a digital optical power meter and photodiode power sensor (Thorlabs PM100D, S120C). Optical power output was measured at the original 12 cm fiber length and then repeatedly after cutting off 1 cm length of fiber to a total of 10 times. This was repeated for three fiber samples of each core-cladding configuration to yield an exponential relation between fiber length and optical power output. The optical loss coefficient was characterized as:

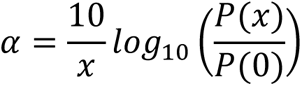

Where *P*(*x*) is the optical power output at length *x*, and *P*(0) s the optical power output at the shortest length measured (∼2 cm).

Numerical apertures of polished, ferrule-coupled waveguides were experimentally measured using the 470 nm fiber-coupled LED light source described above and a low-light CMOS camera (FLIR Blackfly, BFS-U3-200S6M-C). Each fiber was mounted on a micrometer-coupled mechanical translation stage. The profile of the output beam was captured at regular relative distances (*Δ* = 0 mm, 1 mm, 2 mm, 3 mm, 4 mm, 5 mm) between the CMOS sensor and the fiber facet (**Supporting Figure S2a**). Beam profiles were fitted to gaussian curves and the lateral beam extent, *d*, for each relative fiber-sensor distance was calculated as the two-sided width of the fitted beam profile (**Supporting Figure S2b**) at which the maximum signal intensity (*L*_0_) had decreased to *L*_0_ × *e*^−2^. NA values were calculated as the angular beam divergence, or increase in 1/*e*^2^ beam diameter at each *Δ*_*i*_, as

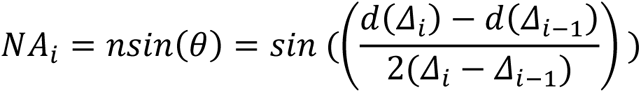

where *n* indicates refractive index of the measuring medium (*n_air_* = 1), and *θ* indicates the angle of beam divergence. For n = 3 waveguides of each composition, final NA values were calculated as linear fits to beam divergence data and compared to theoretical or manufacturer-reported values (**Supporting Figure S2c**). For custom-drawn polymer waveguides, theoretical numerical apertures (NA) were calculated from reported refractive indices of the core (*n_core_*) and cladding (*n_cladding_*) materials as:

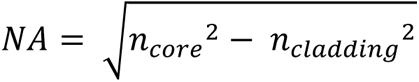

### Agarose brain synthesis and fluorescein injection

A 5-times concentrated (5X) Tris-borate-EDTA (TBE) solution was prepared by dissolving 54 g of Tris(hydroxymethyl)aminomethane (TRIS) (17926, Thermo Scientific) and 27.5 g of boric acid (B0394, Sigma-Aldrich) in 20 ml of 0.5M pH 8 Ethylenediaminetetraacetic acid (EDTA) (AM9260G, Invitrogen) and 900 ml of deionized water. 0.6% w/v Agarose LE (50-192-7938, Fisher Scientific) was dissolved in heated in 1X TBE. 1500ml of the dissolved 0.6% agarose gel was poured into a brain mold (B009S5SL90, Amazon) and allowed to cool overnight at room temperature. Following brain phantom gelation, 10µl of fluorescein (AAL1325122, Fisher Scientific) dissolved in 1X TBE at a concentration of 10 µg/ml was injected into the agarose brain using an extended length pipette tip.

### Electrochemical Characterization

Electrochemical properties of the 20 µm-diameter CNT microelectrodes at the fiber probe tip were evaluated in a three-electrode cell using a potentiostat (Gamry Instruments, Gamry Interface 1010e). A graphite rod was used as the counter electrode, an Ag/AgCl reference electrode (Sigma Aldrich) was used as the reference, and 10 mM phosphate buffered saline (PBS) at pH 7.4 (Fisher Scientific) was used as the electrolyte. Electrochemical impedance spectroscopy (EIS) was performed in a range 1 Hz – 100 kHz with 10 mV_p-p_ AC driving voltage. Cyclic voltammetry (CV) was performed at a sweep rate of 50 mV/s. The water window limits of the CNT electrodes were determined by incrementally increasing the negative limit of the CV scan until water reduction was observed (beginning at −1.5 V), then the positive limit of the CV scan until a linear, resistive behavior indicating water oxidation was observed (beginning at +1.2 V). Cathodic charge storage capacity (CSC_C_) was determined from CV scans from –1.5 to +1.2 V by taking the time integral of the cathodic current. To measure the cathodic charge injection capacity (CIC_C_), voltage transient testing was performed using chronopotentiometry with biphasic, charge-balanced current pulses with t_c_ = t_a_ = 250 μs and t_ip_ = 250 μs for currents ranging from 100 μA to 500 µA. The maximum cathodic potential (E_mc_) was determined as the instantaneous voltage 10 µs after the end of the cathodic current pulse. E_mc_ values were plotted as a function of injected current amplitude, and the linear relation was determined to estimate the current limit at which the electrode would reach its cathodic limit, –1.5 V. CIC_C_ was defined as:

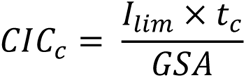

Where *I*_*lim*_ is the cathodic current limit, *t*_*c*_ is the cathodic pulse width, and GSA is the electrode geometric surface area.

### Microfluidic Channel Characterization

To characterize the performance of microfluidic channels in converged POLI fiber samples (n = 3), saline was injected at 25, 50, or 75 nL/min into the microfluidic channel using a syringe pump system (UMP3 Syringe pump and NanoFil syringe, World Precision Instruments). A capillary attached to the microfluidic outlet was visualized under a microscope, and images were continuously recorded with video capture. A small volume (∼1 μL) of mineral oil was withdrawn into the outlet capillary such that the oil-water interface could be visualized. The interface position was tracked during the infusion and used to report flow rate as a function of the known capillary diameter (250 or 375 μm).

### Mechanical Characterization

The stiffness of converged POLI fibers (n = 3, 360-380 × 400-420 μm^2^) compared to a comparably sized silica fiber (400 μm diameter) was characterized with dynamic mechanical analysis (DMA, TA Instruments, Discovery Q850). DMA measurements were performed in single-cantilever mode over a 17.5 mm sample length at 37 °C using 10 μm vertical deflections.

### Dual-site Implantation in Mesolimbic Pathway

All animal procedures were approved by the MIT committee on Animal Care and performed in accordance with the IACUC protocol 0121-002-24. The plasmid pAAV-hSyn-dLight1.1 was purchased from Addgene (#111066) and packaged into AAV9 serotype in-house to a titer of 7.7 x 10^12^ vg/mL (protocol described below). The Cre-dependent red excitatory opsin pAAV-CamKIIa-ChrimsonR-mScarlet-KV2.1 in AAV9 serotype was purchased from Addgene (#124651-AAV9) with a titer of ≥5 x10^12^ vg/mL. Wild type C57BL/6J (The Jackson Laboratory, #000664) and DAT-Ires-Cre (The Jackson Laboratory, #006660) mice aged 8 weeks were used for the study, and were housed in a normal 12 h light/dark cycle with standard chow diet and water ad libitum. All surgeries were performed under aseptic conditions. Mice were anesthetized with 1-2% isoflurane, placed on a heat pad in a stereotaxic head frame (Kopf Instruments), and injected subcutaneously with slow-release buprenorphine (ZooPharm, 1.0 mg kg^−1^). Ophthalmic ointment (Puralube) was applied to the animal’s eyes to retain moisture. A midline incision was performed along the scalp, then the skull was repositioned by aligning and levelling Lambda and Bregma landmarks. Implantation and injection coordinates were established following the Mouse Brain atlas by Paxinos and Franklin as follows: ventral tegmental area (VTA; ML +0.45, AP -3.3, DC -4.3) and the nucleus accumbens (NAc; ML +1.25, AP +1.2, DV -4.3).^[65]^ The dual-fiber device was positioned above the animal’s head and aligned to the stereotaxic frame using a custom-designed 3D-printed holder to enable accurate positioning. The microfluidic channels of each probe were connected to a NanoFil Syringe and UMP3 Microinjection pump (World Precision Instruments) via flexible poly(vinyl alcohol) tubing, and the entire line was primed with sterile phosphate buffered saline, taking care to eliminate any air bubbles. Craniotomies were performed using a rotary tool (Dremel Micro 8050) and a carbon steel burr (Heisinger, 19007-05) at each implantation target, with two additional craniotomies drilled over the contralateral cortex for placement of skull screws to serve as the reference and ground electrodes for electrophysiology recordings. The two stainless steel skull screws (McMaster-Carr #90910A310) attached to the dual-fiber device were fixed to the skull with a T1 torx screwdriver such that the tip of each screw contacted the cortical surface. Solutions of viral vectors were then loaded into the microfluidic channel of each fiber probe as follows: 500 nL of AAV9-dLight1.1 was drawn into the tip of the NAc-targeted fiber and 500 nL of AAV9-ChrimsonR was drawn into the tip of the VTA-targeted fiber. The dual fiber device was then lowered into the brain to the coordinates listed above. Following implantation, viruses were infused from each fiber as follows: 250 nL of virus was injected at a rate of 50 nL/min. The device was then cemented to the skull using 3 layers of C&B-Metabond adhesive acrylic (Parkell) followed by dental cement (Jet Set-4) to cover the base of the device and the skull screws. The mouse was subcutaneously injected with carprofen (5 mg kg^-1^) and sterile Ringer’s solution (0.6 mL) prior to being returned to the home cage, placed partially on a heating pad. Post-implantation, animals were provided with food and water ad libitum, were monitored for 3 days for signs of overall health, and were provided with carprofen injections (0.6 mL, 0.25 mg mL^-1^ in sterile Ringer’s solution) as necessary. For biocompatibility studies, 4-5 month old male DAT-Ires-Cre mice (n=4) were prepared for craniotomies as described above. Implantation coordinates were set to NAc; ML -1.25, AP +1.2, DV -4.3 and NAc; ML +1.25, AP +1.2, DV -4.3 for the POLI and silica fibers respectively. Using a dual device holder, one POLI fiber and one silica fiber were lowered into each mouse following the craniotomy. The device was adhered to the skull as described above, following the same post-implantation protocol.

### Functional In Vivo Experiments

In vivo experiments were performed a minimum of 14 days post-implantation to allow the dLight1.1 indicator and ChrimsonR opsin sufficient time for expression. Animals were anesthetized with 1-2% isoflurane, placed on a heat pad, and ophthalmic ointment (Puralube) was applied to the animal’s eyes. For fiber photometry and optogenetic stimulation experiments, the Neurophotometrics FP3002 system was used, and for electrical stimulation and electrophysiology experiments the Ripple Grapevine Scout with Pico+Stim headstage was used, and these systems were coupled with digital I/O to enable time-syncing. For ChrimsonR optogenetic stimulation trials, a 635 nm laser from the FP3002 system was used to deliver 10 mW/mm^2^ optical power from the tip of the fiber in 10 ms pulses at 5 Hz or 20 Hz frequencies, with 60 s between pulse trains. The laser stimulation parameters were controlled from a MATLAB API script controlling the Ripple Grapevine system, via a BNC digital output to the FP3002 to drive the laser, such that a digital output timestamp was captured in the electrophysiology recordings each time a laser pulse was delivered. Electrophysiology signals were recorded at a sampling rate of 30 kHz. For fiber photometry recordings, the 470 nm (dLight/GCaMP) and 415 nm (isosbestic) LEDs in the FP3002 system were each calibrated to deliver 100 µW of optical power out of the tip of the fiber, and recordings were performed at a 130 Hz sampling rate (65 Hz per wavelength alternating between 470 and 415). The patch cord was photobleached for 1 hour prior to each photometry recording experiment. At the start of each photometry recording, a TTL pulse was sent via BNC digital output to the Ripple Scout system to enable time-syncing between the photometry recordings and the electrophysiology data. In trials where electrical stimulation was delivered during fiber photometry recording, a 5 min baseline photometry signal was recorded prior to initiating stimulation, and TTL pulses were sent from the Ripple Scout system and recorded along with the photometry data to enable time-syncing. Electrical stimulation was applied as biphasic, charge-balanced, cathodic first pulses from a CNT electrode on the fiber, with a skull screw as GND/return. In all experiments with electrical stimulation applied to evoke DA release, the following stimulation parameters were used based on previous literature: 60 Hz, 2 ms/phase biphasic, 200 µA, 0.5 s stimulation trains, 30 s interval between stimulation trains.^[66,67]^ In the experiments probing DA dynamics before and after cocaine exposure, fiber photometry recordings of dLight1.1 were performed while electrical stimulation pulses were delivered to the VTA or NAc (only one region stimulated per animal). Baseline photometry signal was recorded for 5 mins before stimulation was initiated. Five stimulation pulse trains (60 Hz, 200 µA, 2 ms/phase biphasic, 0.5 s stimulation trains, 60 s interval) were delivered, then stimulation was paused and an intraperitoneal dose of cocaine (20 mg/kg) solution was injected. 5 minutes after the IP injection of cocaine, electrical stimulation was restarted using the same parameters and 5 trains of stimulation were delivered.

### Fiber photometry in vivo

Ten Thy1-GcaMP6s mice (The Jackson Laboratory #024275, 8-weeks old) were implanted with either 400 µm silica waveguide (0.5 NA, Thorlabs; FP400URT, n = 6 mice), or with 200 µm or 400 µm PMMA/THVP waveguide (n = 1 mouse, and n = 3 mice respectively). Using the same surgical approach as outlined above, each animal was implanted with an optical waveguide targeting their somatosensory barrel cortex (S1BF; ML ±3mm, AP - 1.2mm, DV -0.4mm). Following a week-long recovery, the animals were anesthetized using a ketamine/xylazine cocktail (ketamine, 100mg/kg; xylazine, 10mg/kg). Fiber photometry was performed using the same experiment approach as outlined above. Calcium signals were expressed as z-score. Following 2 minutes of baseline recording, whiskers contralateral to implanted S1BF were mechanically stimulating using a cotton tipped applicators, following a brushing motion. This stimulation was repeated every 30 seconds for a total of 7 times.

### Data Analysis

Electrophysiology and fiber photometry data were analyzed in MATLAB. Fiber photometry data were processed as follows: Data were low-pass filtered below 25 Hz using a 2^nd^ order Butterworth filter with zero phase distortion.^[7]^ The first 2 min of the baseline recording were discarded to eliminate the effects of fast photobleaching that occurs at the start of the recording. The 470 nm and 415 nm signals were fit with a biexponential decay function to approximate the photobleaching dynamics, and the fitted photobleaching curves were subtracted from each signal to produce 470 nm and 415 nm signals that were corrected for photobleaching. The bleach-corrected 415 nm signal was regressed onto the bleach-corrected 470 nm signal with robust non-negative linear regression, thus scaling the isosbestic signal onto the 470 nm signal. This scaled isosbestic signal was then subtracted from the 470 nm bleach-corrected signal, producing a signal that has been corrected for both photobleaching and motion artifacts as recorded by the 415 nm isosbestic. Baseline fluorescence F0 was then calculated as the mean fluorescence value of the corrected 470 nm signa, during the baseline period. ΔF/F was finally calculated as ΔF/F = (corrected 470 nm signal – F0) / F0. Electrophysiology data were filtered into the spikeband from 300 Hz – 6 kHz using a 2^nd^ order Butterworth bandpass filter with zero phase distortion. To extract and sort neuron spike units, spike waveforms were detected by threshold crossings, the waveforms were extracted and aligned by the spike peak, principal components analysis (PCA) was applied to the extracted waveforms, and k-means clustering was applied to cluster the spike waveforms and sort them into different neuronal units.

### AAV Packaging

AAV9-hSyn-dLight1.1 was produced following a previously reported protocol.^[55]^ Briefly, pAAV2/9n (Addgene #112865), pAAV-hSyn-dLight1.1 (Addgene #111066), and pHelper (CELL BIOLABS, INC.) were used for PEI-mediated triple transfection of HEK293T cells. The culture medium was collected at 72 h and 120 h, and the cells were harvested at 120 h post-transfection, followed by purification by ultracentrifugation. The AAVs were collected in Dulbecco’s phosphate-buffered saline (DPBS) (Gibco) with 0.001% Pluronic F-68 (Gibco). AAV titers were determined using a Taraka Bio AAV real-time PCR titration kit (#6233). pAAV2/9n was a gift from James M. Wilson (Addgene plasmid #112865; http://n2t.net/addgene:112865; RRID:Addgene_112865). pAAV-hSyn-dLight1.1 was a gift from Lin Tian (Addgene plasmid #111066; http://n2t.net/addgene:111066; RRID:Addgene_111066).

### Histology

Wild-type or DAT::Cre mice were implanted in the VTA and NAc POLI fibers or commercial silica waveguides (300 µm, FT300UMT Thorlabs) were anesthetized with isoflurane, injected intraperitoneally with Fatal-Plus (100 mg kg−1), and transcardially perfused with 50 ml of ice-cold PBS followed by 50 ml of ice-cold 4% paraformaldehyde (PFA) in PBS. The devices were carefully explanted, and the brains were removed and additionally fixed in 4% PFA in PBS for 24 h at 4 °C, then stored in PBS afterwards. Coronal slices (40-µm thickness) were prepared using a vibratome (Leica, VT1000S) and a razor blade (Electron Microscopy Sciences, 72002) in ice-cold PBS. The slices were then stored in PBS at 4 °C in the dark until staining. Slices were permeabilized with 0.3% v/v Triton X-100 and blocked with 3% bovine serum albumin in PBS for 30 min. Slices were incubated overnight at 4 °C in the blocking solution with primary antibody (Iba1: rabbit anti-Iba1, ab178846 Abcam, 1:200 dilution; GFAP: Goat anti-GFAP, ab53554 Abcam, 1:200 dilution; CD68: mouse anti-CD68, ab31630, Abcam, 1:200 dilution; GFP: rabbit anti-GFP, A-11122 Invitrogen 1:200 dilution). Following incubation, slices were washed three times with PBS. The slices were then incubated with a secondary antibody (Donkey anti-Goat Alexa Fluor 555, A32816, Thermofisher, 1:1,000 dilution; Goat Anti-Rabbit Alexa fluor 488, ab150077, Abcam, 1:1,000 dilution; Goat Anti-Mouse Alexa Fluor 647, ab150115, Abcam, 1:1,000 dilution) for 1 h at room temperature on a shaker followed by an additional three washes with PBS. Slices were then incubated with DAPI (4′6-diamidino-2-phenylindole) (1:20,000) for another 20 min, and washed three times with PBS. Fluoromount-G (SouthernBiotech) was used for mounting slices onto glass microscope slides (48311-703, VWR) and covered with No. 1.5 coverslips (CLS-1764-2250, Erie Scientific). A white light laser scanning confocal microscope (Stellaris 5, Leica) was used for imaging with a 10X objective. Regions of interest were chosen based on the implant locations. Aivia was used to quantify the immune response by calculating the area over which the signal of the immune markers was present then normalizing to the image acquisition area. Intensity measurements were calculated using pixel-based total intensity measurements of the regions where signal was detected.

### Surgery for MRI

All animal procedures were conducted in accordance with National Institutes of Health guidelines and with the approval of the MIT Committee on Animal Care with the with the IACUC protocol number 0721-059-24. The MRI experiments were performed with female Sprague-Dawley rats, age 8–12 weeks (Charles River Laboratories, Wilmington, MA). Eight rats were used for in vivo imaging experiments.

For stereotactic surgery, rats were anesthetized with isoflurane (3% for induction, 1.5% for maintenance) and placed on a water heating pad from Braintree Scientific (Braintree, MA) to keep body temperature at 37 °C. After stereotaxic frame fixation and topical lidocaine application a 3 cm lateral incision extending from bregma to lambda was made to expose the skull. Craniotomies (0.5 mm) were drilled unilaterally over the right Nucleus accumbens area (NAc), 6.0 mm anterior 1.7 mm and 3.5 mm lateral to bregma. After 30 minutes, POLI fiber implant or a standard setup with custom-made silver bipolar electrode (two silver wires with each: inner diameter: 0.127 mm; outer diameter 0.178 mm, twisted together, A-M Systems, # 786000) and a 200 µm fiber optic cannula (CFMXC10, Thorlabs) was lowered to 7.5 mm below the surface of the skull through the craniotomy and held in place by the stereotactic arm. The implants were fixed to the skull with dental cement (Secure Resin Cement, Parkell). Rats were allowed to recover for 8 weeks until the start of imaging experiments.

### MRI imaging and stimulation

Rats were anesthetized with isoflurane (induction 3% and maintenance 0.5% via nose-cone) and dexmedetomidine (0.05 mg/mL) for imaging. High field MRI acquisition was performed with a 20 cm-bore 9.4 T Bruker small animal scanner. A custom-made 30 mm single surface coil was used as a transceiver. Field inhomogeneity was minimized via MAPSHIM protocol in Paravision 6.0.1 software. Anatomical scans were acquired using a *T*_2_-weighted rapid acquisition with refocused echoes (RARE) pulse sequence with 18 slices of 1 mm thickness, 20 mm × 20 mm field of view (FOV), image size 200 × 200, echo time (*TE*) 34.7 ms, repetition time (*TR*) 2 s, and 8 averages. Functional scans were performed using T2*-weighted EPI sequence for detection of stimulus-induced BOLD contrast, with following parameters: *TE* of 16 ms, *TR* of 2 s, *FOV* 20 mm × 20 mm, image size 40 × 40, 18 slices with slice thickness of 1 mm.

Deep brain stimulation (DBS) of 0.1mA for 2s at a frequency of 60Hz was applied preceded by 10s baseline scan and followed by 48s resting scan combining into 1 cycle. 30 cycles were applied under continuous functional EPI scans.

### MRI Data Processing and Analysis

Images were reconstructed using the ParaVision 6.1 software and further processed by the National Institute of Health AFNI software package. Functional imaging time series were preprocessed in following steps: Slice timing correction, motion correction by a least-squares rigid-body volume registration algorithm, voxel-wise intensity normalization, spatial smoothing with Gaussian spatial kernel of 0.5 mm full-width at half-maximum and spatial resampling, all steps were performed in AFNI. High-resolution anatomical images of each animal were registered to a Waxholm coordinate space rat brain atlas. Functional images were aligned to anatomical images. After averaging 30 cycles of DBS, maximum signal amplitude during an interval of 6s after stimulation onset was determined and compared to averaged signal during the preceding baseline interval via z-test to evaluate the statistical significance of z-scores based on maximum signal amplitude minus average baseline divided by the standard deviation of the baseline. A Student’s t-test was used to evaluate statistical significance of z-scores. To count the number of voxels in each individual with signal loss due to the implants, a threshold of SNR< 5 was applied in the slice where the implant was visible on the anatomical images in the right hemisphere.

### FSCV Chemicals and Materials

Sodium chloride, sodium sulfate, calcium chloride, sodium phosphate monohydrate, and paraformaldehyde were purchased from Sigma-Aldrich (St. Louis, MO). Potassium chloride and magnesium chloride were purchased from Thermo Fisher Scientific (Waltham, WA). Dopamine (DA) hydrochloride was dissolved in 0.1 M HClO_4_ to create a 10 mM DA stock solution. Phosphate buffered saline (PBS; 131.25 mM NaCl, 3.00 mM KCl, 10 mM NaH_2_PO_4_, 1.2 mM MgCl_2_, 2.0 mM Na_2_SO_4_, and 1.2 mM CaCl_2_) was used at pH 7.4 to dilute the DA stock solution to 1.0 μM. Deionized water (EMD Millipore, Billerica, MA) was used to prepare all aqueous solutions.

### FSCV and electrochemical measurements

Fibers were conditioned by surface polishing using a KT Brown micro-pipette beveller (model BV-10, Sutter Instruments, Novato, CA). Before each use, each fiber was inserted into a glass capillary (A&M Systems, Inc., Carlsborg, WA) previously pulled using a vertical PE-22 Electrode Puller (Narishige, Tokyo, Japan) to a diameter of 500 µm, exposing 1 mm of the entire fiber length. The capillary was used to backfill with 1M KCl and attach the fiber, ensuring a proper connection with the electrode holder (Warner Instruments, Holliston, MA). The tips were sealed by using 5-min epoxy (J-B weld, Sulphur Springs, TX), and the silver wire in the electrode holder was connected through the backfilled capillary. All solutions were injected into a flow cell from a syringe at a rate of 2 mL min^-1^ by a pump (Harvard Apparatus, Holliston, MA). The injection and analyte-buffer mixing was facilitated with a six-port, stainless steel air actuator (VICI Valco Instruments, Houston, TX). DA voltammograms were obtained using a Chem-Clamp potentiostat (Dagan Corp., Minneapolis, MN) coupled to a UNC breakout box (UNC Electronics Shop, Chapel Hill, NC) with a 1 MΩ head stage. HDCV software (UNC at Chapel Hill) was used for data acquisition and analysis. Ag/AgCl wires were used as the reference electrodes. A 3 kHz low-pass filter was used to detect DA with a triangular waveform that scanned from −0.4 to 1.3 V and back at 100 V/s at 10 Hz.

### FSCV experiments in vivo

All animal experiments were performed as approved by the Animal Care and Use Committee (ACUC) of the University of Virginia. Male Sprague Dawley rats (Charles River Laboratories, Wilmington, MA, USA) between 280 - 320 g were anesthetized with urethane (0.3 ml/l00 g, 5% saline solution., i.p.). A local anesthetic (bupivacaine) was used on exposed skin and muscle tissue during surgery. The rat was placed in a stereotaxic frame, and craniotomies were drilled precisely to place the stimulating and working electrodes according to the atlas of Paxinos and Watson.^[51]^. The fiber was lowered into the NAc core (+1.3 mm AP, +2.0 mm ML, −7.1 mm DV), and a bipolar stimulating electrode (Plastics One, Roanoke, VA, USA) was lowered into the VTA (−4.7 mm AP, +0.9 mm ML, −8.5 mm DV). DA was detected using FSCV at the NAc following stimulation in the VTA with 24 biphasic electrical pulses (2 ms, 300 µA, 60 Hz) delivered every 5 minutes. The dorsoventral coordinate of the electrodes in the VTA was adjusted to maximize the DA release. DA release in the NAc core was for 30 minutes. The recording electrodes were then subject to post-experiment calibration. Following in vivo measurements, a solution of 2.0 µM dopamine in PBS was used to recalibrate the linear relationship used to correlate peak current values detected in vivo to DA concentration.

## Supporting Information

Supporting Information is available from the Wiley Online Library or from the author.

## Acknowledgements

N.D., M.-J.A, T.M.C., and P.M. contributed equally to this work. This work was supported in part by Lore Harp McGovern Addiction Initiative at the McGovern Institute for Brain Research at MIT, National Institute of Neurological Disorders and Stroke (R01-NS115025-01A1, P.A.; R01-NS121014 B.J.V, R01-NS125663 B.J.V.), Pioneer Award from the National Institutes of Health and National Institute for Complementary and Integrative Health (DP1-AT011991, PA), and the K. Lisa Yang Brain-Body Center at MIT. P.M. is a recipient of a MathWorks Engineering Fellowship. A.B.S. was a recipient of the Lore Harp McGovern Fellowship.

## Conflict of Interest

N.D., M.-J. A. and P.A. are co-founders and have a financial interest in NeuroBionics Inc.

Received: ((will be filled in by the editorial staff))

Revised: ((will be filled in by the editorial staff))

Published online: ((will be filled in by the editorial staff))

