## Supporting Information for "Fiber-based Probes for Electrophysiology, Photometry, Optical and Electrical Stimulation, Drug Delivery, and Fast-Scan Cyclic Voltammetry In Vivo"

Supplementary Note 1. Notch Fiber.

During the design and fabrication of the POLI fiber, we fabricated a functionally equivalent fiber from the same components, termed the Notch fiber, using post-draw integration of carbon nanotube (CNT) yarn rather than convergence during the fiber thermal drawing process. This was done initially to test the concept in a cost-effective way, since convergence integration of six CNT microwires into the final POLI fiber required >60 m of costly 20 μm-diameter CNT microwire. Fabrication of the notch fiber was nearly identical to that of the final POLI fiber, including the same materials for the optical waveguide, poly(methyl methacrylate) (PMMA) core and THVP (terpolymer of tetrafluoroethylene, hexafluoropropylene, and vinylidene fluoride) cladding, with the same dimensions, however notched grooves along three sides of the preform were drilled in place of the six hollow channels found in the POLI fiber preform. These notches were filled with SEBS elastomer (styrene ethylene butylene styrene) to retain their shape during the drawing process. After the notch fiber was drawn, these strips of SEBS (final width of ~20 μm) were peeled from the notches on the sides of fiber sections, and the CNT microwire was pressed into these three groves along the sides of the fiber. The fiber was then coated with 5 μm of Parylene-C via chemical vapor deposition to electrically insulate and fix the CNT microwires in place. This produced a final fiber device that was functionally equivalent to the POLI fiber, but with three CNT electrodes rather than six.

**Supporting Table 1.** Glass transition temperature ($T_g$) and refractive index values of polymers used in the POLI fiber. PMMA: poly(methyl methacrylate), PC: polycarbonate, COC: cyclic-olefin-copolymer, THVP: terpolymer of tetrafluoroethylene, hexafluoropropylene, and vinylidene fluoride, SEBS: styrene-ethylene-butylene-styrene.

| Material | Refractive index, *n* (wavelength) | Glass transition temperature $T_g$ (°C) |
|---|---|---|
| PMMA | 1.4956 (488 nm) [1] | 105-120 [2] |
| PC | 1.5976 (488 nm) [1] | 130 - 170 [3] |
| COC | 1.543 (488 nm) [4] | 158 [5] |
| THVP | 1.35 (589 nm) [6] | 130 (Tm) [6] |
| SEBS | N/A | 90 [7] |

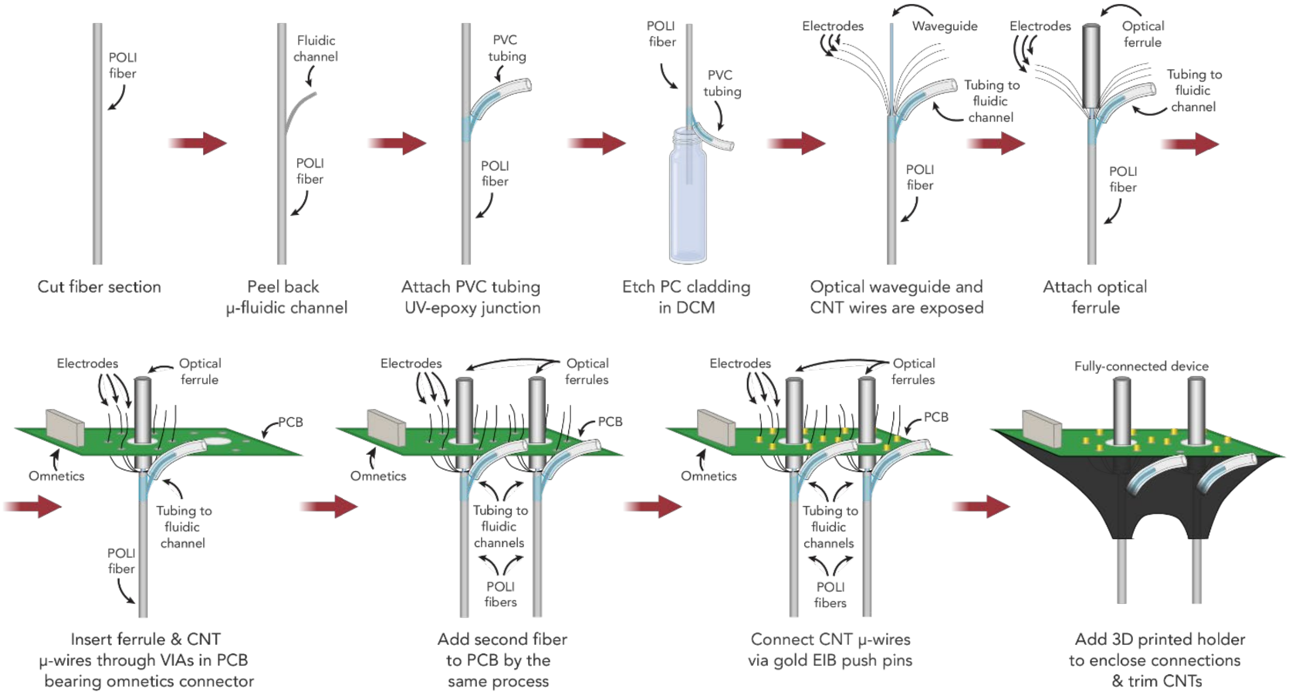


**Supporting Figure S1.** Diagram for POLI fiber interfacing with the multifunctional backend.

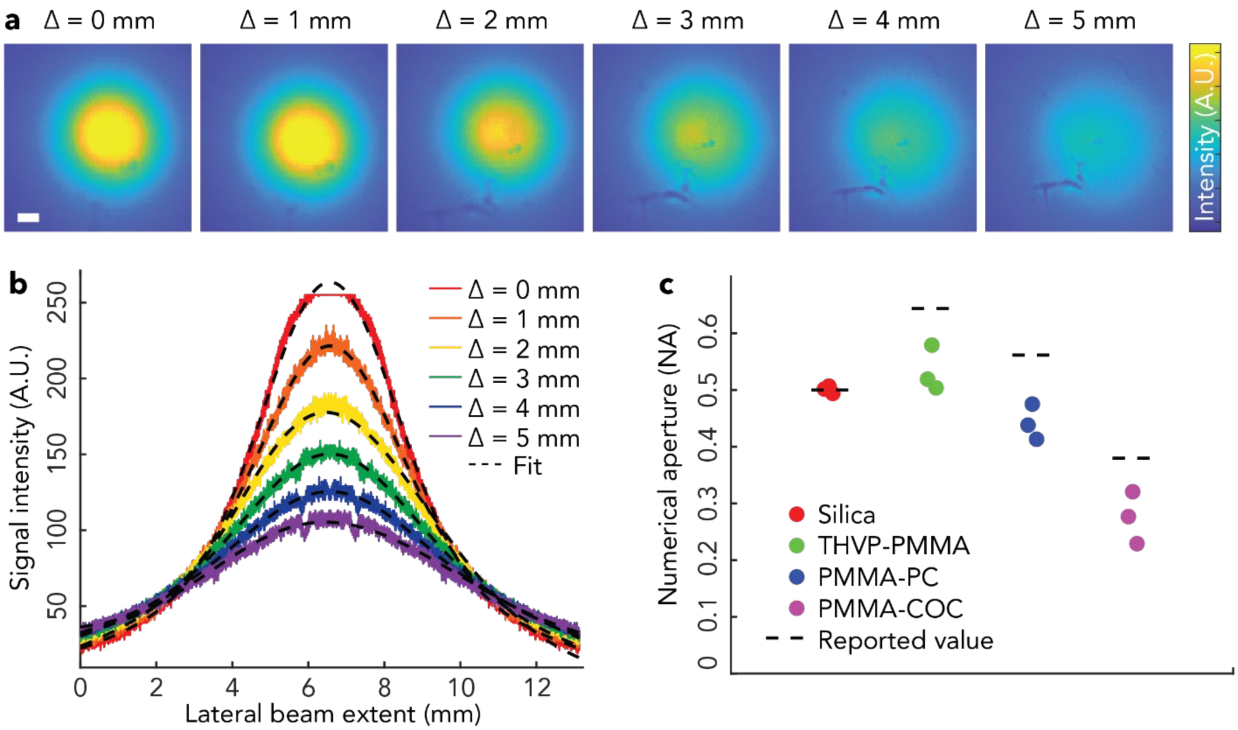


**Supporting Figure S2. Quantification of numerical aperture (NA). (a)** Representative beam profile images of PMMA-PC waveguide output light distribution at increasing distances (delta) between the fiber tip and the camera sensor (scale bar = 1 mm). **(b)** Fitted gaussian beam profiles for PMMA-PC waveguide at each delta. **(c)** NA quantification of each polymer waveguide (N = 3 fibers) based on beam dispersion method and comparison to values calculated from reported refractive indices (THVP-PMMA, PMMA-PC, PMMA-COC) or reported by manufacturer (silica). PMMA: poly(methyl methacrylate), PC: polycarbonate, COC: cyclic-olefin-copolymer, THVP: terpolymer of tetrafluoroethylene, hexafluoropropylene, and vinylidene fluoride, SEBS: styrene-ethylene-butylene-styrene.

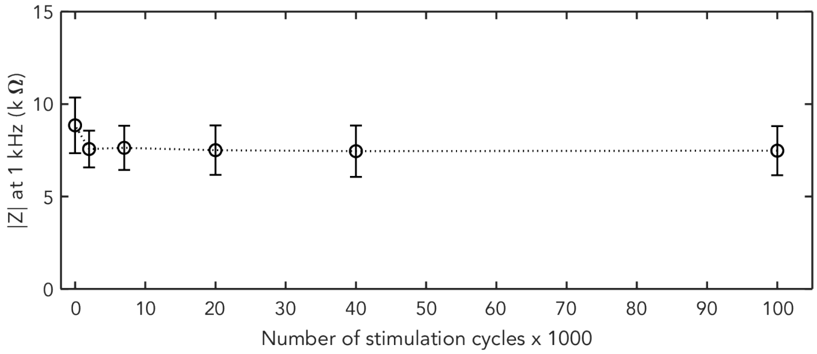


**Supporting Figure S3. Stability of POLI fiber-integrated CNT electrodes for electrical stimulation.** Mean impedance magnitude at the 1 kHz reference frequency of 20 µm CNT electrodes subjected to 100,000 cycles of stimulation pulsing with the following parameters: 200 µA, 50 Hz, biphasic, cathodic-first, 200 µs phases with 25 µs interphase interval (N=5 electrodes, 100,000 pulses). A slight decrease in impedance is observed after the first 2000 cycles of stimulation due to the expected electrode conditioning effect, beyond which the impedance is stable.

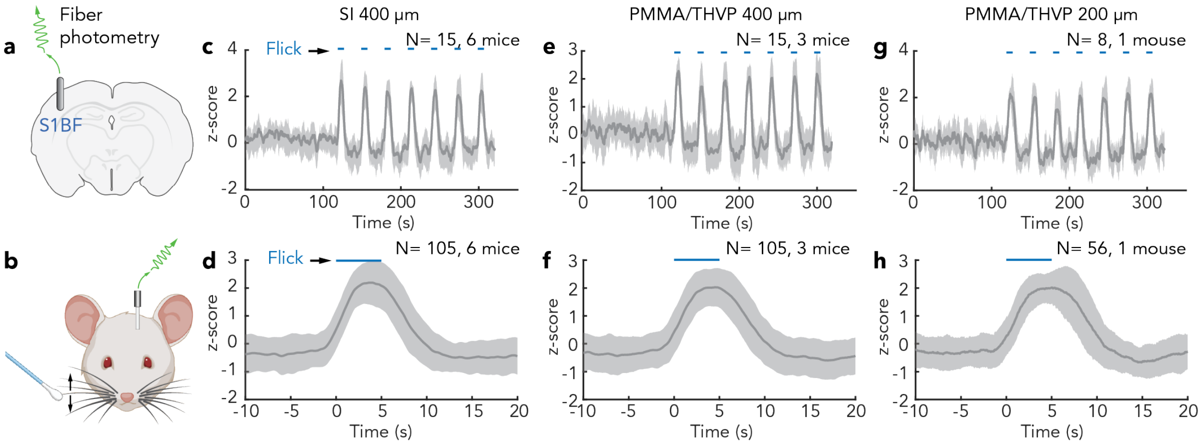


**Supporting Figure S4.** Validation of fiber photometry in vivo. **(a)** Schematic of the implantation site. Eight-weeks old Thy1-GCaMP6s mice were implanted with a 400 µm silica waveguide (n = 15 trials, 6 mice - 3 females and 3 males), 400 µm PMMA/THVP waveguide (n = 15 trials, 3 mice - 2 male and 1 female), or 200 µm PMMA/THVP waveguide (n = 8 trials, 1 female mouse). All implants targeted S1BF area. **(b)** Schematic of the experiment. The whiskers contralateral to the implantation site were mechanically stimulated with a brush, while neuronal activity was recorded via GCaMP6s photometry. **(c-h)** Average GCaMP6s signal recorded via a 400 µm silica waveguide (c-d), 400 µm PMMA/THVP waveguide (e-f), or a 200 µm (g,h) during the whisker stimulation experiment (c,e,g), or represented as a stimulation onset aligned response (d,f,h); each blue tick corresponds to a whisker brush. Data is represented as mean ± s.e.m.

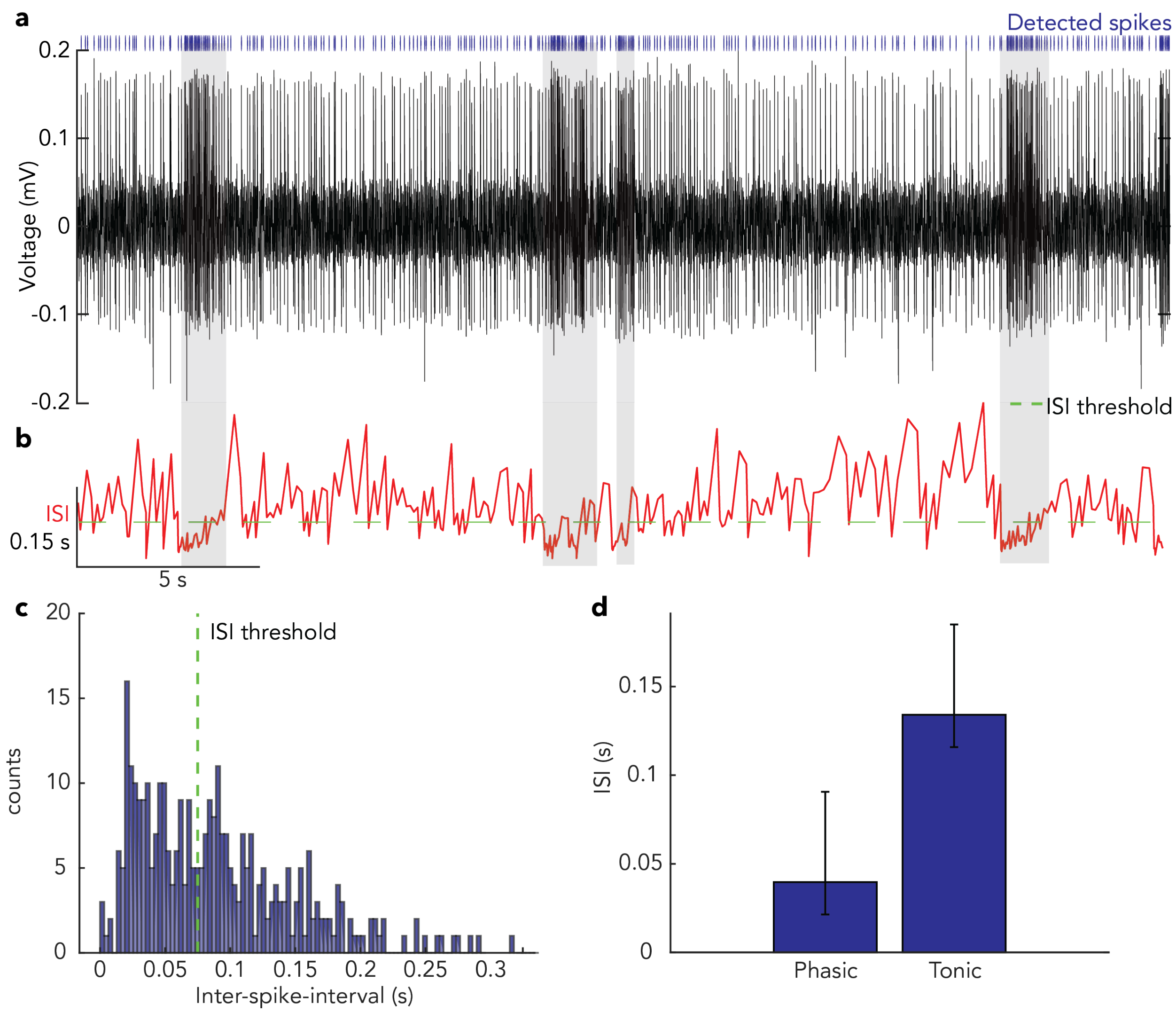


**Supporting Figure S5. Quantification of endogenous spiking activity in the VTA**. **(a)** Raw electrophysiological recording also shown in **Figure 4b**. **(b)** The inter-spike interval (ISI) for the electrophysiological data in (a) was quantified throughout periods of tonic and phasic electrophysiological activity. **(c)** An ISI threshold of 0.075 s (13 Hz) was chosen based on visual assessment to differentiate phasic and tonic activity (b). **(d)** Phasic activity below this threshold was found to have an average ISI of 0.040 s (25 Hz), and tonic activity above this threshold was found to have an average ISI of 0.13 s (7.5 Hz).

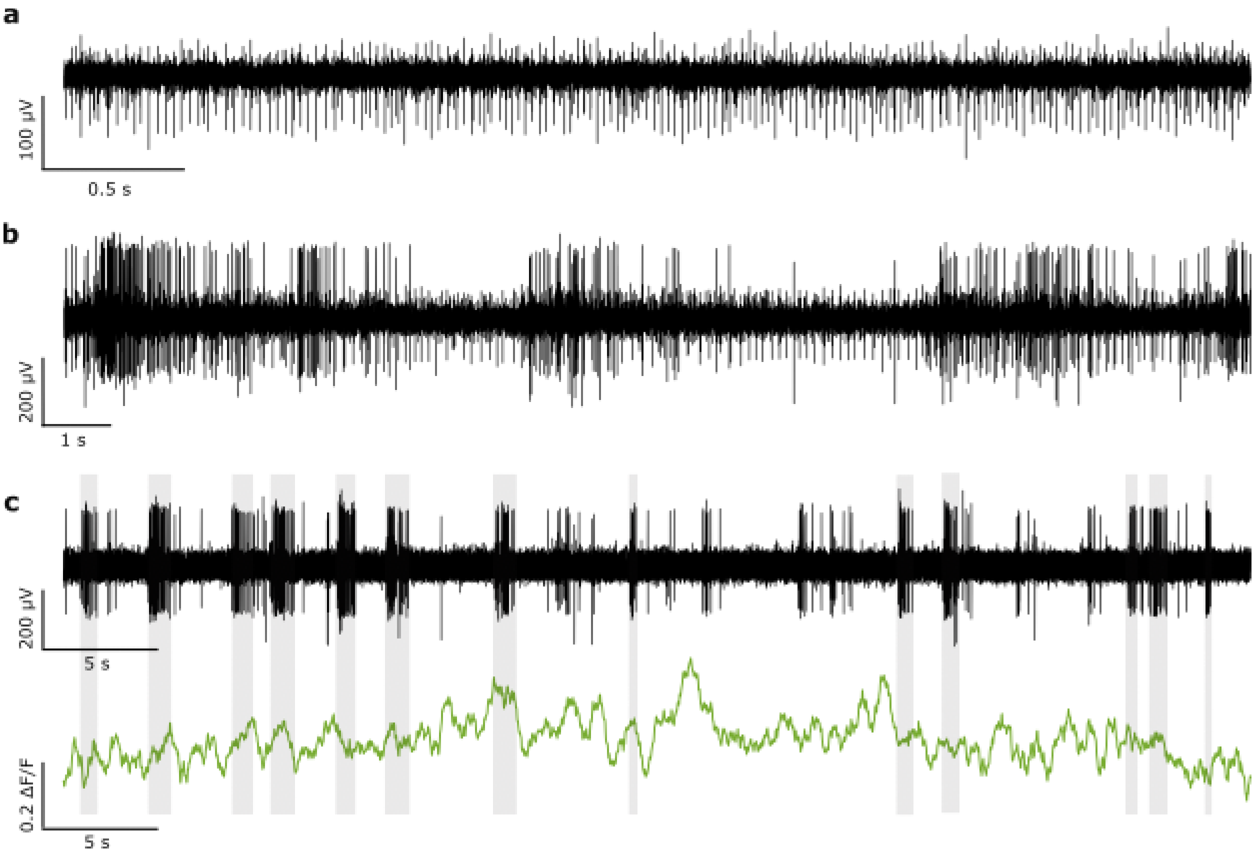


**Supporting Figure S6. Endogenous firing of VTA neurons and corresponding DA transients recording in NAc**. **(a)** Raw electrophysiological recording of endogenous firing activity in the VTA. Phasic firing was not observed in this animal. **(b)** Raw electrophysiological recording of endogenous firing activity in the VTA of a different animal, this time with notable phasic activity. **(c)** Time-aligned electrophysiological recording in the VTA with photometric recording of dLight1.1 fluorescent transients in the NAc of a third animal. Epochs of bursting are highlighted in grey.

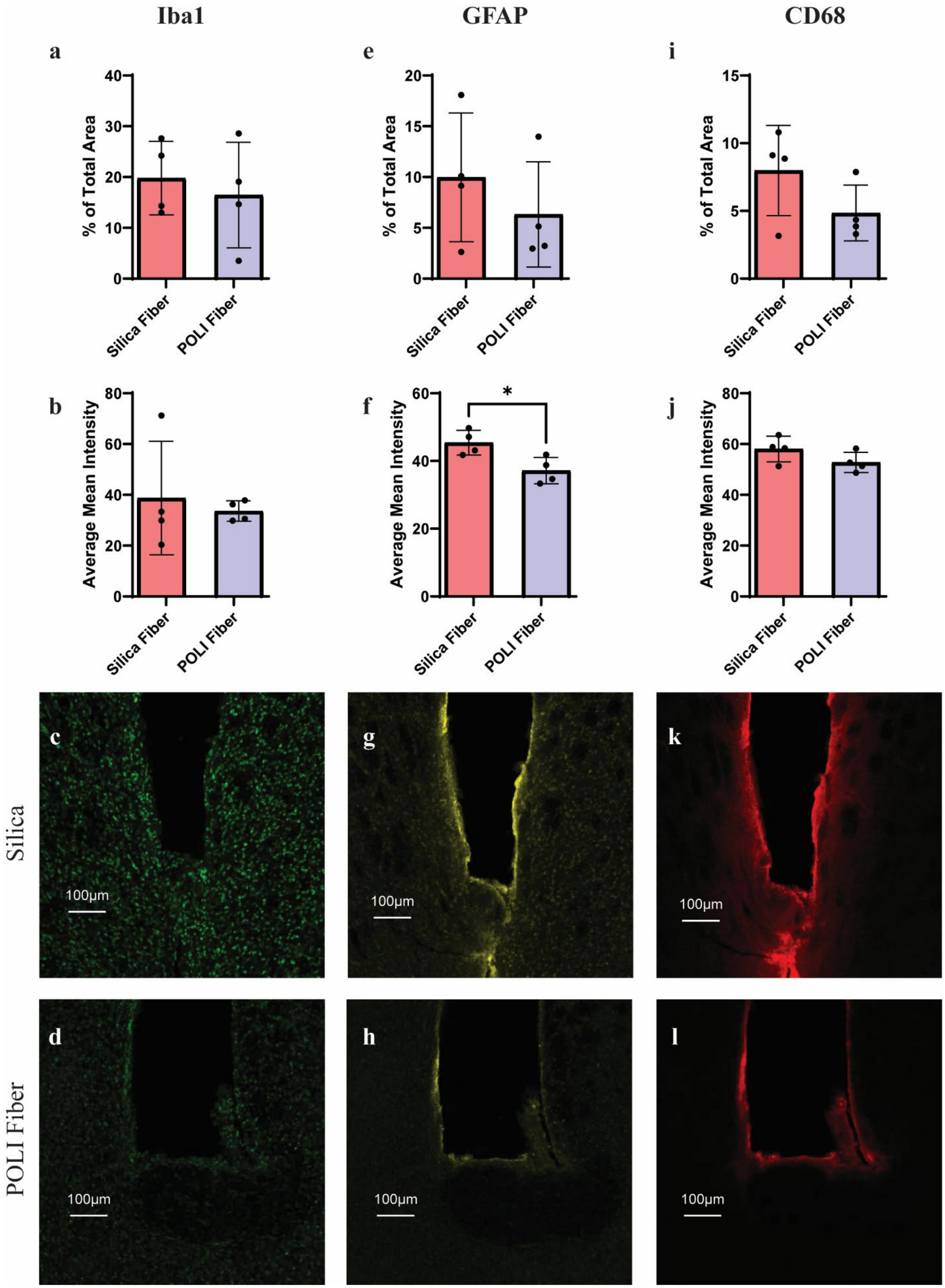
Iba1
GFAP
CD68
a
b
c
d
e
f
g
h
i
j
k
l
% of Total Area
Average Mean Intensity
Silica Fiber
POLI Fiber
*
Silica
POLI Fiber
100µm

**Supporting Figure S7. Immunohistochemical evaluation of chronically bilaterally implanted silica and POLI fibers.** Immunofluorescent quantification of astrocytic (GFAP) and microglial (Iba1, CD68) markers surrounding a 300-400μm section of POLI-fiber and a 400 μm silica fiber implanted into the NAc of male DAT$^{IREScre}$ mice (N = 4) (HC PLAPO CS2 10x/0.40 Dry objective, WLL: 85% Power, Speed: 400Hz, 405: Intensity (3.80), Gain (33.76); 499: Intensity (0.5), Gain (27.62); 554: Intensity (3.66), Gain (25.49), 653: Intensity (2), Gain (33.76), scale bar = 100 μm). Average fluorescent area and average fluorescence intensity of Iba1 (a-d), GFAP (e-h), and CD68 (i-l) as well as representative confocal images at the implant tips of POLI fiber (-1.25ML, +1.2AP, -4.3DV) and silica fiber (+1.25ML, +1.2AP, -4.3DV) 1 month post implantations. GFAP average mean intensity; p = 0.0421, Student's t-test. Data are presented as mean values +/- s.d.
